# RNA G-quadruplexes forming scaffolds for α-synuclein aggregation lead to progressive neurodegeneration

**DOI:** 10.1101/2023.07.10.548322

**Authors:** Kazuya Matsuo, Sefan Asamitsu, Kohei Maeda, Kosuke Kawakubo, Ginji Komiya, Kenta Kudo, Yusuke Sakai, Karin Hori, Susumu Ikenoshita, Shingo Usuki, Shiori Funahashi, Yasushi Kawata, Tomohiro Mizobata, Norifumi Shioda, Yasushi Yabuki

## Abstract

Synucleinopathies, including Parkinson’s disease, dementia with Lewy bodies, and multiple system atrophy, are triggered by the aggregation of α-synuclein, leading to progressive neurodegeneration^1,2,3,4,5,6,7,8^. However, the intracellular mechanism of α-synuclein aggregation remains unclear. Here we show that assembly of RNA G-quadruplexes forming scaffolds for α-synuclein aggregation, contributing to neurodegeneration. Purified α-synuclein binds RNA G-quadruplexes directly through the N-terminus. RNA G-quadruplex itself undergoes phase separation and assembly by Ca^2+^, accelerating the sol–gel phase transition of α-synuclein. In α-synuclein preformed fibrils-treated neurons, RNA G-quadruplexes assembly composed of synaptic mRNAs co-aggregates with α-synuclein upon Ca^2+^ excess influx into cytoplasm, eliciting synaptic dysfunction. Forced assembly of RNA G-quadruplexes using an optogenetic approach evokes α-synuclein aggregation, neuronal dysfunction and neurodegeneration. Administration of 5-aminolevulinic acid, a prodrug of protoporphyrin IX that prevents phase separation of RNA G-quadruplexes^9^, attenuating α-synuclein aggregation, neurodegeneration, and progressive motor deficits in α-synuclein preformed fibrils-injected synucleinopathy mice. Together, assembly of RNA G-quadruplexes due to dysregulation of intracellular Ca^2+^ homeostasis accelerates α-synuclein phase transition and aggregation may contribute to pathogenesis of synucleinopathies.

## Main

α-synuclein (αSyn) is an intrinsically disordered protein (IDP) of 140 amino acids^10^. A relatively high concentration of αSyn is present in synaptic vesicle membranes; it is also present as a natively unfolded monomer in the cytoplasm^11^. Although the physiological role of αSyn remains elusive, it is presumably involved in synaptic vesicle clustering, docking, fusion, and recycling^12^. In pathological conditions, monomeric αSyn assembles into toxic oligomers, protofibrils, and amyloid-like fibrils in synucleinopathies, such as Parkinson’s disease (PD), dementia with Lewy bodies (DLB), and multiple system atrophy (MSA)^1,2,3,4,5,6^. The aggregates composed of αSyn, which are known as Lewy bodies (LBs), are a neuropathological hallmark of synucleinopathies, and their pathogenesis has been attributed to neuronal loss^1–3, 7, 8^. However, the mechanism underlying the transformation of native αSyn into pathological aggregates is not completely understood.

Liquid–liquid phase separation (LLPS) represents a ubiquitous phenomenon in which molecules transition from a homogeneous state into a dense phase with different physiochemical properties (e.g., molecular crowding), inducing the formation of liquid droplets, hydrogels, and aggregates^13–17^. Proteins and RNAs often undergo LLPS due to multivalent macromolecular interactions with themselves and other biomolecules^9, 13, 18^. LLPS results in the formation of membraneless ribonucleoprotein (RNP) organelles enriched in RNA and RNA-binding proteins (RBPs), including nucleoli, Cajal bodies, nuclear speckles, stress granules (SGs), processing bodies (P-bodies), and neuronal RNA granules^19, 20^. The liquid–solid phase transition via LLPS could be an important intermediate step for aggregate formation in neurodegenerative disease-associated IDPs, such as αSyn^21^, tau ^22^, fused in sarcoma (FUS)^23^, TAR DNA-binding Protein of 43 kDa (TDP-43)^24^, and heterogeneous nuclear ribonucleoprotein A1 (hnRNP A1)^25^, whereas the intrinsic factor that facilitates the phase transition has not been identified.

We found that RNA G-quadruplexes (rG4s), which are quadruple-stranded structures formed by contiguous guanines^26^, bind to αSyn and induce aggregation. rG4 itself undergoes LLPS in a Ca^2+^-dependent manner, acting as a scaffold for αSyn aggregation. In αSyn pre-formed fibrils (PFFs)-treated mouse neurons forming LB-like inclusions similar to human synucleinopathies^27, 28^, αSyn co-aggregated with rG4 assembly consisting of synaptic mRNAs associated with intracellular Ca^2+^ overload, thereby demonstrating synaptic dysfunction. In pharmacological experiments, oral administration of 5-aminolevulinic acid (5-ALA), which intracellularly generates protoporphyrin IX (PPIX) that inhibits rG4s LLPS^9, 29^, suppressed αSyn aggregation and improved behavioral motor deficits in αSyn PFF-treated mice. These results provide evidence that rG4s assembly induces sol–gel phase transition of αSyn, eliciting progressive neurodegeneration.

### *α*Syn binds to rG4s and undergoes sol–gel phase transition

First, we analyzed the temporal changes in the aggregation of endogenous αSyn using primary mouse cortical neurons treated with human αSyn PFF. In untreated controls, αSyn was found to be intracellularly diffuse assembling at synaptophysin-positive presynapses along microtubule-associated protein 2 (MAP2)-positive neurites (**Extended Data Fig. 1a**). However, after PFF treatment, αSyn showed significantly higher inhomogeneity in the cytoplasm, existing as granules (**Fig. 1a, b**). The immunoreactivity of αSyn phosphorylated at serine 129 (pS129), a central feature of the LB inclusion, increased in a time-dependent manner and was evident eight days after PFF treatment (**Fig. 1b** and **Extended Data Fig. 1b**). Of note, αSyn granules mainly co-localized with decapping mRNA 1A (Dcp1a)-positive P-bodies (43.0%) but rarely with fragile X mental retardation protein (FMRP)-positive RNA granules (3.2%) or p62-positive aggresomes (18.9%) four days after the treatment (**Fig. 1a, b** and **Extended Data Fig. 1c**). However, αSyn granules localized primarily in the aggresomes (43.9%) but rarely in the P-bodies (6.1%) or the RNA granules (3.1%) eight days after the treatment (**Fig. 1a, b** and **Extended Data Fig. 1c**). Since human antigen R (HuR)-positive SGs were not detected upon PFF treatment (**Extended Data Fig. 1c**), neurons were treated with sodium arsenite for SG formation. Although both HuR-positive SGs as well as αSyn granules were detected in the cytoplasm after the treatment, they did not co-localize (**Extended Data Fig. 1d**).

**Fig. 1:**
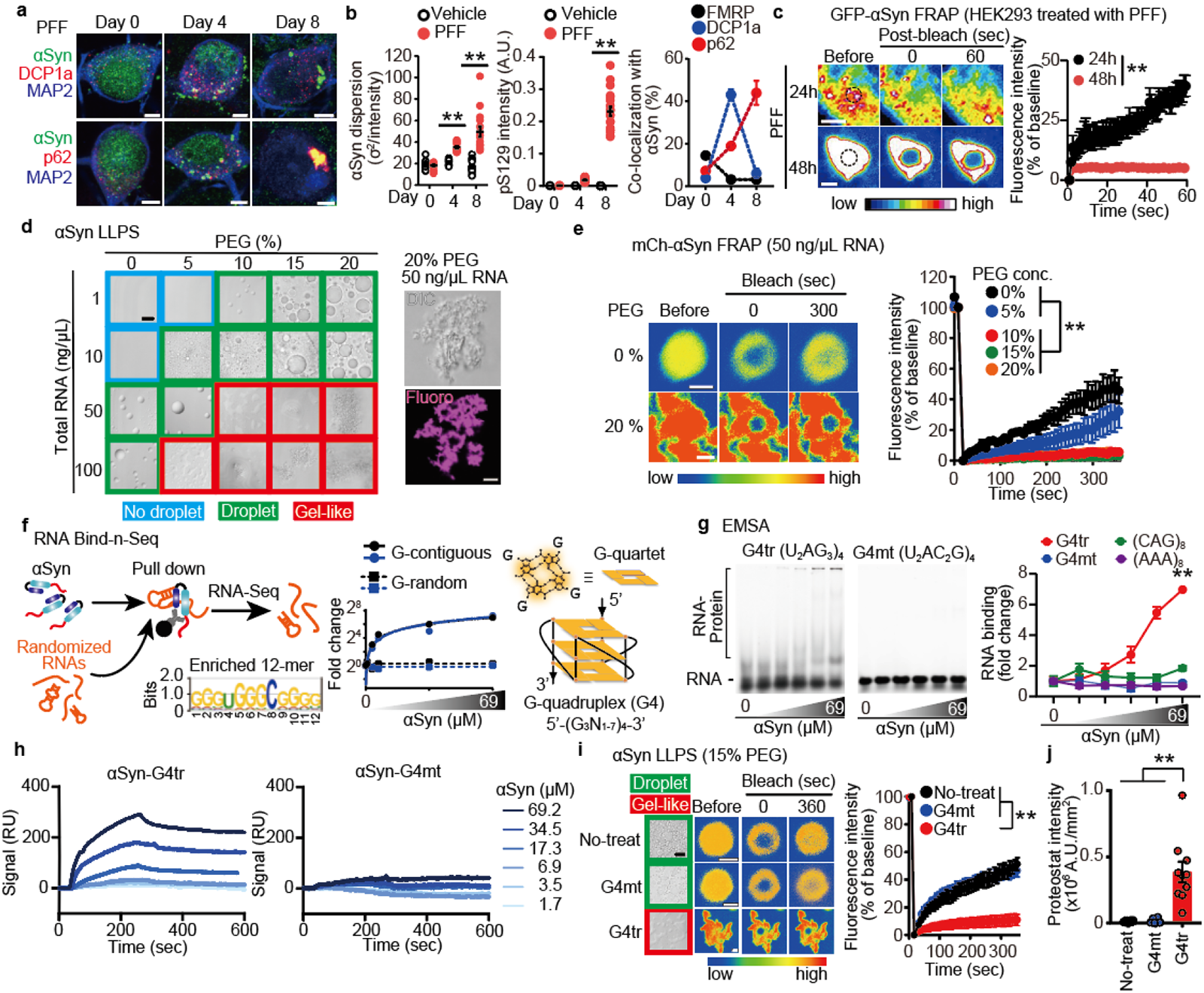
RNA G-quadruplexes initiate *α*-synuclein (*α*Syn) sol–gel phase transition. **a**, Representative images of MAP2 (blue) and mouse αSyn (green) with DCP1a or p62 (red) in mouse cultured neurons following pre-formed fibril (PFF) treatment. Scale bars, 5 μm. **b**, Analyses of αSyn dispersion (left), pS129^+^ immunoreactivity relative to MAP2^+^ area (center), and co-localization of αSyn with markers of RNA granule and aggresome (right) in mouse cultured neurons after PFF treatment (images shown in **Extended Data Fig. 1b, c**). **c**, αSyn fluorescence recovery after photobleaching (FRAP) assay of PFF-treated HEK293T cells following transient expression of GFP-αSyn. Scale bars, 2 μm. **d**,**e**, *In vitro* αSyn phase separation (d) and FRAP assay (e) dependent on polyethylene glycol (PEG) and total RNA. Scale bars, 5 and 1 μm, respectively. DIC, differential interference contrast. **f**, Schematic of RNA Bind-n-Seq experiment for αSyn and the top enriched RNA motif (left). Fold enrichment of the top two 12-mers (circle marks) and two randomly chosen 12-mers (square marks) across αSyn concentrations (0, 0.69, 3.45, 6.9, 34.5, and 69 μM; center). Schematic illustration of intramolecular parallel G4 (right). **g**, electrophoresis mobility shift assay for the interaction of αSyn with RNA oligomers in the presence of 100 mM NaCl. αSyn concentrations were the same as those of RNA Bind-n-Seq (**f**). **h**, surface plasmon resonance sensorgrams for the interaction of αSyn with G4tr or G4mt. RU, response unit. **i**, *In vitro* αSyn phase separation (left) and FRAP assay (right) with G4tr or G4mt in the presence of 15% PEG. Scale bars, 5 and 1 μm, respectively. **j**, Proteostat intensity within phase-separated RNA/αSyn in the presence of 15% PEG. Data are presented as mean ± standard error of mean. \*\**P* < 0.01 by two-way (**b,c**,**e**,**g** and **i**) and one-way (**j**) Analysis of variance with Bonferroni’s multiple comparisons test. Number of replicates is shown in Supplementary Table 9.

To assess the dynamic nature of αSyn with PFF treatment in cells, we utilized HEK293 cells overexpressing green fluorescent protein (GFP)-tagged αSyn in the present study. The GFP-tagged αSyn granules exhibited rapid recovery kinetics 24 h after PFF treatment in fluorescence recovery after photobleaching (FRAP) assay (**Fig. 1c**, **Extended Data Fig. 1e** and **Supplementary Video**), and the GFP-tagged αSyn granules were not co-localized with pS129 or p62 (**Extended Data Fig. 1f**). However, the fluorescence recovery was markedly reduced and αSyn granules co-localized with pS129 and p62 48 h after the treatment (**Fig. 1c**, **Extended Data Fig. 1e, f** and **Supplementary Video**). These results suggest that αSyn transiently forms granules with RNPs via LLPS, subsequently forming aggregates contributing to the pathogenesis of synucleinopathies. Thereafter, we employed clustered regularly interspaced short palindromic repeats/Cas9 technology in several human cell lines (HEK293, Hela, and Hutu-80 cells) for endogenous tagging with mNeonGreen at *SNCA* loci to reveal the dynamic nature of endogenous αSyn with PFF treatment; however, both the N- and C-terminal tagging was unsuccessful owing to unexplained cell death.

Since our data suggested that RNP granules containing RNAs and/or proteins provide scaffolds for initial αSyn aggregation, we subsequently examined whether the addition of cell-derived RNAs or proteins undergo sol–gel phase transition of αSyn *in vitro*. Purified αSyn (69 μM) underwent LLPS according to the concentration of polyethylene glycol (PEG), a molecular crowding agent, whereas no droplet formation was observed without PEG (**Extended Data Fig. 2a**). Thereafter, we examined whether RNAs impact αSyn LLPS by adding total RNAs extracted from Neuro-2a cells. RNA addition decreased the PEG concentration required for αSyn LLPS and elicited a transition from a droplet to a gel-like state (**Fig. 1d**). Fluorescence imaging studies using mCherry-tagged αSyn (mCh-αSyn) confirmed that these gel-like aggregates contain αSyn (**Fig. 1d**). In FRAP experiments in the presence of RNAs, the fluorescence recovery of αSyn was faster in the droplets but not in the gel-like states (**Fig. 1e**). Similar experiments were performed with nuclease-treated proteins: αSyn droplets formed in the presence of PEG; however, no transition to a gel-like state occurred (**Extended Data Fig. 2b**).

These observations motivated us to examine the nucleotide-binding capacity of αSyn and its sequence; therefore, we subsequently performed RNA Bind-n-Seq^30^ using recombinant human αSyn and a pool of random 24-nt RNA oligonucleotides *in vitro* (**Fig. 1f**). The top 10 αSyn binding RNA motifs were significantly guanine-rich sequences (**Extended Data Fig. 3a–e** and **Supplementary Table 1**). Several runs at different concentrations of αSyn revealed that these motifs require guanines in a contiguous but not random sequence, thereby suggesting rG4s as a potential structure for αSyn binding (**Fig. 1f** and **Extended Data Fig. 3f, g**). To investigate whether αSyn preferentially interacts with rG4s, we performed an electrophoresis mobility shift assay (EMSA) using various RNA oligonucleotide structures: G4tr—a typical rG4, telomeric repeat-containing RNA (TERRA) (UUAGGG)_4_ repeats, G4mt—a TERRA mutant that is unable to form rG4 (UUACCG)_4_ repeats, a hairpin structure—(CAG)_8_ repeats, and polyA— single-stranded and non-structured (AAA)_8_ repeats (**Extended Data Fig. 4a**). αSyn demonstrated high affinity only for G4tr and little or no affinity for other RNA structures (**Fig. 1g** and **Extended Data Fig. 4b**). In quantitative surface plasmon resonance (SPR) binding analysis, αSyn demonstrated much higher selectivity for G4tr (*K_D_* = 3.63 ± 0.63 μM) compared to that with G4mt (*K_D_* = n.d.) (**Fig. 1h** and **Extended Data Fig. 4c, d**). αSyn is defined by three regions: 1) the N-terminal region (Nterm; residues 1–60), which is basic and composed of seven imperfect repeats of 11 amino acids that contain a 4-residue core with the consensus sequence KTKEGV; 2) the non-amyloid-β component region (NAC; residues 61–95), which is hydrophobic and forms the core of the fiber during aggregation; and 3) the C-terminus (Cterm; residues 96–140), which is a highly negatively charged region that binds to metal ions^31^. To identify the domains responsible for rG4-binding capability, we constructed three mutant αSyns: ΔNterm, ΔNAC, and ΔCterm. SPR analysis demonstrated ΔNAC and ΔCterm but not ΔNterm segments showing binding affinity for G4tr, suggesting that the N-terminus of αSyn preferentially binds to rG4s (**Extended Data Fig. 4c, d**). We also obtained similar results using the Quartz Crystal Microbalance system.

As for αSyn LLPS *in vitro*, G4mt demonstrated no additional effects on αSyn droplet formation in the presence of 15% PEG, while G4tr evoked αSyn transition into gel-like states (**Fig. 1i**). Furthermore, αSyn ΔNterm mutant formed droplets but not gel-like aggregates with G4tr (**Extended Data Fig. 4e**). In FRAP studies, the fluorescence recovery of αSyn droplets was rapid when formed alone, with G4mt, and as ΔNterm with G4tr, whereas that of gel-like assemblies with G4tr was not recovered (**Fig. 1i** and **Extended Data Fig. 4e**). The intensity of Proteostat signals was significantly higher in gel-like assemblies of αSyn with G4tr compared to that in droplets of αSyn alone, αSyn with G4mt, and ΔNterm with G4tr (**Fig. 1j** and **Extended Data Fig. 4f**). Considered together, rG4s bind directly to the N-terminus of αSyn, accelerating the sol–gel phase transition *in vitro*.

### Ca^2+^-induced rG4 assembly is essential for *α*Syn aggregation

We examined the intracellular mechanism underlying the rG4s-induced αSyn aggregation on pathological conditions. rG4 itself has the propensity to undergo LLPS *in vitro*, which can be boosted electrostatically by Mg^2+^ divalent cation^16, 18^. Indeed, G4tr underwent LLPS without PEG; however, G4mt did not (**Extended Data Fig. 5a**). Moreover, other divalent cation Ca^2+^ promoted G4tr LLPS droplets and enlarged the size of droplets (**Extended Data Fig. 5a,b**). Furthermore, the Ca^2+^-induced increase in droplet size was markedly suppressed by the addition of EGTA, a Ca^2+^ chelator (**Extended Data Fig. 5b**). Notably, G4tr droplets co-aggregated with αSyn in the presence of Ca^2+^ (**Fig. 2a**). In the absence of G4tr, αSyn droplets or gel-like aggregates were not observed regardless of the presence of Ca^2+^ (**Fig. 2a**). Ca^2+^ transitioned G4tr-αSyn coacervates to a gel-like state thereby significantly reducing FRAP, whose effect was abolished by EGTA (**Fig. 2b**). As expected, Proteostat signals were detected only in the G4tr-αSyn complex in the presence of Ca^2+^ (**Fig. 2c**). Moreover, lower concentration of αSyn (34.5 μM) underwent sol–gel transition in the presence of G4tr and Ca^2+^ (**Extended Data Fig. 5c**). These results suggested that Ca^2+^-induced rG4 assembly is essential for the efficient initiation of αSyn aggregation.

**Fig. 2:**
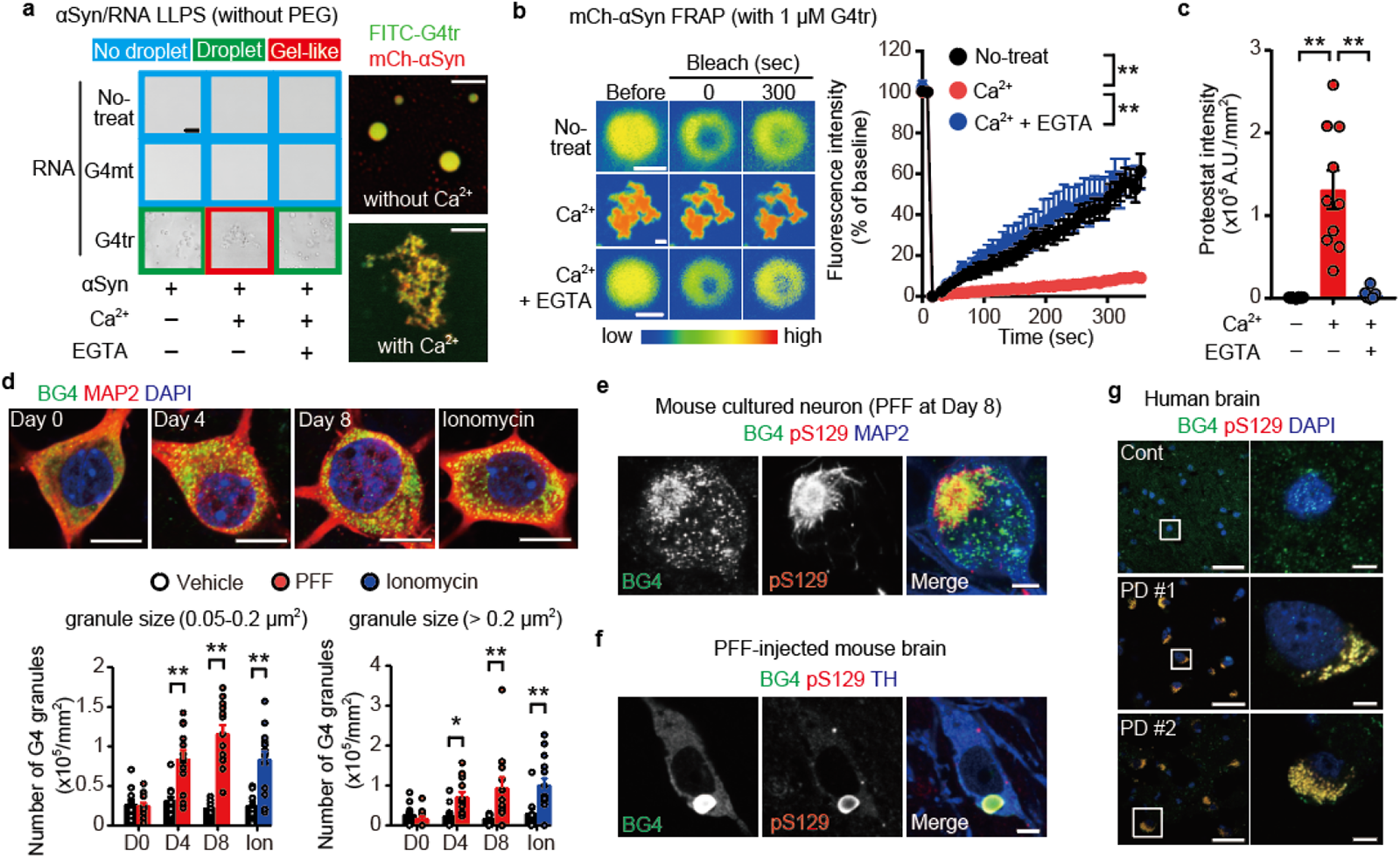
Ca^2+^ triggers RNA G-quadruplexes -induced *α*-synuclein (*α*Syn) sol–gel phase transition. **a**, Representative images of *in vitro* αSyn phase transition dependent on G4tr and Ca^2+^. Scale bars, 10 (left) and 5 (right) μm, respectively. Veh., vehicle. **b**, *In vitro* αSyn fluorescence recovery after photobleaching assay dependent on G4tr and Ca^2+^. Scale bars, 1 μm. **c**, Proteostat intensity within phase-transitioned RNA/αSyn dependent on Ca^2+^. **d**, Representative images (top) and quantification of the number (bottom) of BG4 granules (green) in primary cultured mouse neurons treated with pre-formed fibrils (PFF) or Ionomycin (Ion; 30 min). Scale bars, 10 μm. **e–g**, Representative images of BG4 (green) and pS129 (red) with MAP2, TH, or 4’,6-diamidino-2-phenylindole (blue) in PFF-treated mouse cultured neurons (**e**), PFF-injected mouse brain (**f**), and human normal control and postmortem brains of patients with Parkinson’s disease (**g**). Low and high magnification scale bars, 20 and 5 μm, respectively. Data are presented as mean ± standard error of mean. \*\**P* < 0.01 by two-way (**b**,**d** (vehicle vs. PFF)) and one-way (**c**) Analysis of variance with Bonferroni’s multiple comparisons test and by two-sided, unpaired Student’s *t*-test (**d** (vehicle vs. Ionomycin)). Number of replicates is shown in Supplementary Table 9.

The persistent and excessive Ca^2+^ entry in neurons by certain metabolic stress conditions could be a major factor in the development of neurodegeneration, including PD^32, 33^. Indeed, in response to PFF-induced cellular stress, cultured neurons demonstrated an excessively elevated intracellular Ca^2+^ concentration (**Extended Data Fig. 5d**)^34^. Thus, we examined whether Ca^2+^-induced rG4 assembly is observed in neurons by immunostaining with G4-specific antibody BG4^35^. PFF and Ca^2+^ ionophore (ionomycin) treatments significantly increased the number of cytosolic G4 granules of both small (0.05–0.2 μm^2^) and large (> 0.2 μm^2^) sizes compared to vehicle-treated controls (**Fig. 2d**). Furthermore, the number of G4 granules co-localized with αSyn significantly increased after PFF treatment (**Extended Data Fig. 6a**). Eight days after PFF treatment, pS129-positive aggregates were extensively co-localized with G4 granules in mouse cultured neurons (**Fig. 2e**). In the synucleinopathy model mice injected with PFF into the dorsal striatum, pS129-positive inclusions were observed in the tyrosine hydroxylase (TH)-positive dopaminergic (DA) neurons of the substantia nigra, which co-localized with large G4 granules (**Fig. 2f**). In the brainstems of postmortem human patients with PD, the majority (> 90%) of pS129-positive inclusions co-localized with large G4 granules (**Fig. 2g** and **Extended Data Fig. 6b**). The immunoreactivity of cytosolic G4 granules seen in asymptomatic controls was completely eliminated by RNase pretreatment, whereas immunoreactivity of cytosolic G4 granules co-localized with pS129-positive inclusions in PD brains was not (**Extended Data Fig. 6b**).

### *α*Syn co-accumulated with rG4 containing synaptic mRNAs following Ca^2+^ overload

To identify the rG4s as scaffolds of αSyn aggregation in neurons, BG4-bound RNAs were analyzed in PFF-treated cultured mouse neurons using RNA immunoprecipitation coupled with sequencing (BG4 RIP-seq)^18^.

We identified 254 and 524 gene transcripts enriched in vehicle- and PFF-treated neurons, respectively, of which 162 were common (cutoff: *P-*value < 0.05 and fold-change > 1.5) (**Fig. 3a** and **Supplementary Tables 2 and 3**). We compared fragments per kilobase of exon per million mapped reads (FPKM) obtained from BG4-enriched fractions with those from IgG-treated fractions to identify the rG4s. To predict quadruplex-forming guanine-rich sequences (QGRS) mapper^36^, more than 95% of BG4-enriched mRNAs were found to have putative rG4-forming sequences with two or three G-tetrad layers (**Fig. 3b, Extended Data Fig. 7a** and **Supplementary Table 4,5**). In the gene ontology (GO) analysis, BG4-binding RNAs obtained from PFF-treated neurons contributed to neurodegeneration (**Extended Data Fig. 7b** and **Supplementary Tables 6 and 7**), and most of these RNAs were linked to the synapse (**Fig. 3c** and **Supplementary Table 8**). Among the rG4 enriched in PFF-treated neurons, we focused on two mRNAs encoding synaptic proteins, *Camk2a* and *dlg4*, which have been validated to fold into G4 structures by biophysical assays^37^ and conserved in both mice and humans (**Extended Data Fig. 7c**). Moreover, these synaptic proteins (i.e., CaMKIIα and PSD95) are essential for synaptic plasticity, an important cellular mechanism underlying learning and memory^38, 39^. We also validated that both mRNAs contain rG4-forming sequences (G4ck and G4ps; **Fig. 3d** and **Extended Data Fig. 7d**). Both G4ck and G4ps co-aggregated with αSyn in the presence of Ca^2+^ *in vitro*, which were detected by Proteostat (**Fig. 3e** and **Extended Data Fig. 7e**). To obtain an insight into the interaction of αSyn with these mRNAs in the context of pathogenesis, we performed *in situ* hybridization together with immunostaining in mouse cultured cortical neurons eight days after PFF treatment. We observed high co-localization of these mRNAs with pS129-positive aggregates in the cytoplasm (**Fig. 3f** and **Extended Data Fig. 7f**).

**Fig. 3:**
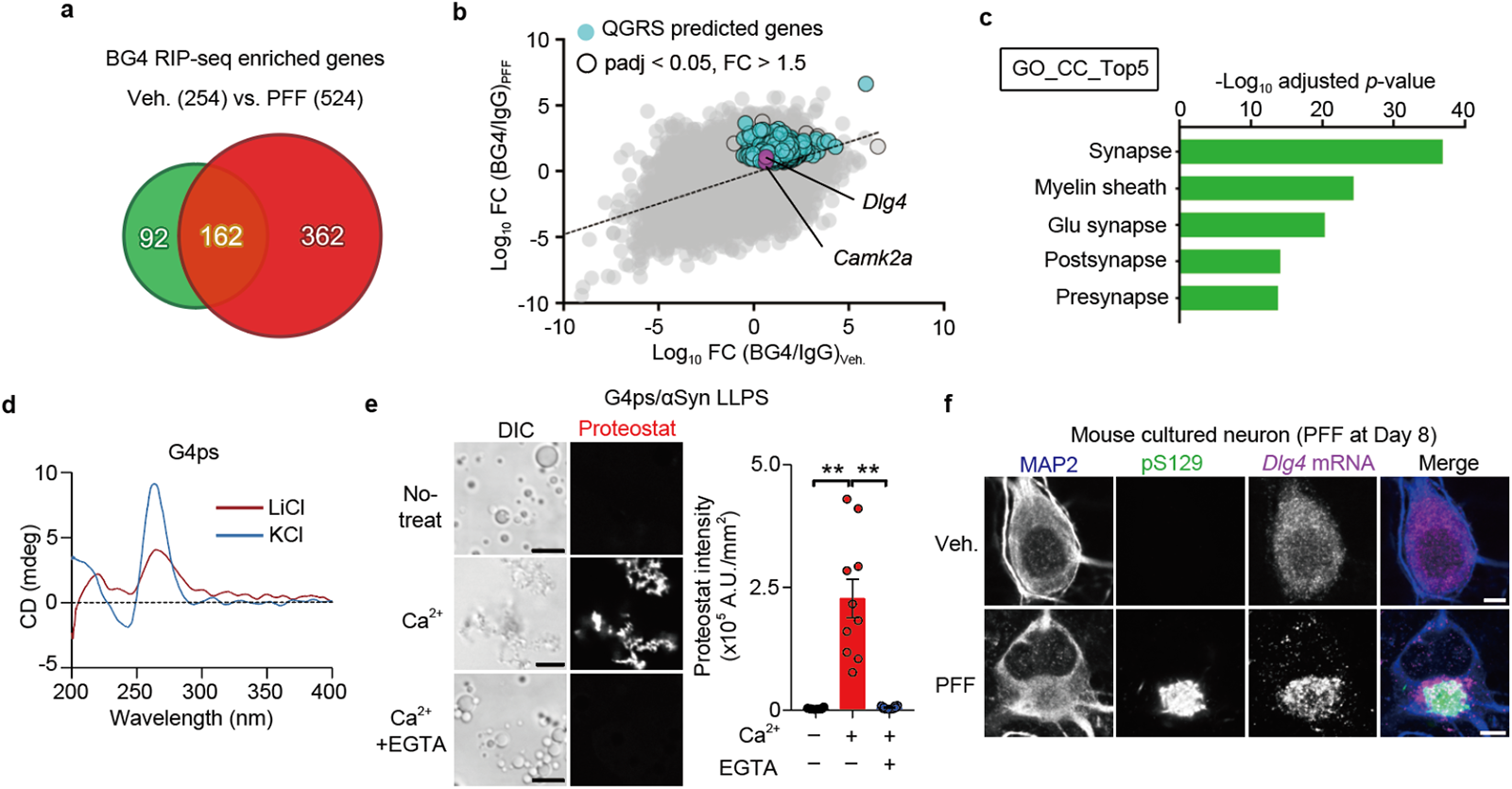
RNA G-quadruplexes-containing synaptic mRNA are involved in *α*-synuclein (*α*Syn) aggregation. **a**, Venn diagram of BG4-enriched RNAs in vehicle- and pre-formed fibrils (PFF)-treated mouse cultured neurons compared with immunoglobulin G (IgG) control. **b**, Scatter plots of BG4-enriched RNAs in vehicle versus PFF treatment. Open black dots show RNAs with *P* < 0.05 and fold-change (FC) > 1.5, and light blue dots exhibit predicted genes in the quadruplex-forming guanine-rich sequences (QGRS) mapper. **c**, Gene Ontology (GO) enrichment analysis by cellular component (CC) terms in BG4-enriched RNAs only detected in PFF-treated mouse cultured neurons. **d**, Circular dichroism (CD) spectra of mouse G4ps (sequence shown in **Extended Data Fig. 9c**) in the presence of KCl or LiCl. **e**, Representative images (left) and quantification (right) of Proteostat intensity within phase-transitioned G4ps/αSyn dependent on Ca^2+^. Scale bars, 5 μm. **f**, Representative confocal images of MAP2 (blue), pS129 (green), and *Dlg4* RNA (magenta) in mouse cultured neurons following PFF treatment. Scale bars, 5 μm. Data are presented as mean ± standard error of mean. \*\**P* < 0.01 by one-way analysis of variance with Bonferroni’s multiple comparisons test. Number of replicates is shown in Supplementary Table 9.

### Optogenetics-induced rG4 assembly evokes *α*Syn aggregation and neuronal dysfunction

To provide direct evidence that rG4 assembly induces αSyn aggregation, we developed an optogenetics-induced rG4 assembly approach, “optoG4 system” under spatiotemporal control of blue light (BL) stimulation. The optoG4 system consists of two plasmids: *G4tr*-*MS2* RNA—rG4-forming (TTAGGG)_23_ repeats tagged 12×*MS2*-hairpin loops^40^ and MCP-Cry2—*MS2*-coat protein tagged mCh-Cry2olig, which functions by co-expressing them in the cells (**Fig. 4a** and **Extended Data Fig. 8a**). We confirmed that MCP-Cry2 clusters were formed in the optoG4-expressing Neuro-2a cells in a BL exposed-time dependent manner (**Fig. 4b**). In the cells co-expressing optoG4 and αSyn, MCP-Cry2 clusters co-assembled with αSyn and *G4tr*-*MS2* RNA 3-h following BL stimulation (**Fig. 4c**). The BL-induced αSyn granules were p62-, pS129-, and BG4-positive (**Extended Data Fig. 8b, c**). Since Cry2olig possesses the property of reversible homo-oligomerization in response to BL stimulation^41^, we examined whether BL-induced αSyn aggregation in the optoG4 system is irreversible. In the cells co-expressing optoG4 and αSyn, MCP-Cry2 formed irreversible complexes containing αSyn and *G4tr*-*MS2* RNA following 3-h BL stimulation and subsequent 3-h withdrawal (**Extended Data Fig. 8d, e**). However, MCP-Cry2 clusters were significantly more dispersed in cells expressing only optoG4 than in cells co-expressing optoG4 and αSyn under the same stimulatory condition (**Extended Data Fig. 8d, e**). We developed “optoG4mt system” using *G4mt*-*MS2* RNA consisting of rG4-unforming (TTACCG)_23_ repeats tagged 12×*MS2*-hairpin loops as a negative control for rG4-forming structure (**Extended Data Fig. 8a, b**). In cells co-expressing optoG4mt and αSyn, MCP-Cry2 and *G4mt*-*MS2* RNA complexes formed 3-h after BL stimulation; however, they did not co-assemble with αSyn or G4s (**Extended Data Fig. 8f**).

**Fig. 4:**
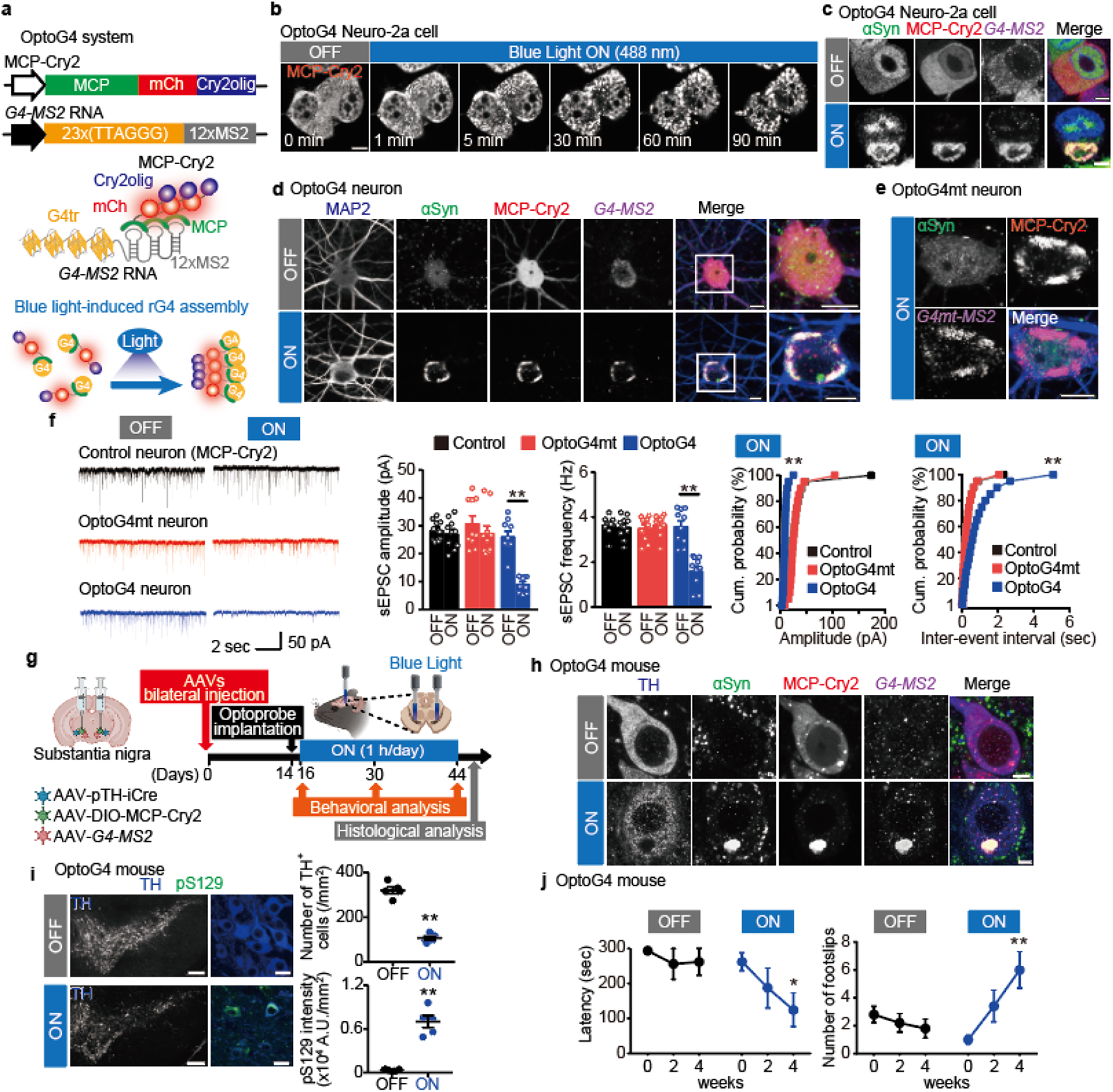
OptoG4-induced *α*-synuclein (*α*Syn) aggregation triggers neurodegeneration. **a**, Schematic of the light-inducible RNA G-quadruplexes assembly approach using the Cry2olig photoreceptor, MCP protein, and 12×*MS2*-hairpin loops. **b**, Representative time-lapse images of Neuro-2a cells challenged with blue light (BL) for the indicated time under expression of optoG4. Scale bar, 20 μm. **c**, Representative images of αSyn (green), MCP-Cry2 (red), *G4*-*MS2* RNA (magenta), and 4’,6-diamidino-2-phenylindole (blue) in optoG4 and αSyn co-expressing Neuro-2a cells with or without exposure to BL stimulation. Scale bars, 5 μm. **d**,**e**, Representative images of MAP2 (blue), αSyn (green), MCP-Cry2 (red), and *MS2* RNA fluorescent *in situ* hybridization (magenta) in optoG4 (**d**) and optoG4mt (**e**) mouse cultured neurons exposed to BL stimulation or darkness. Scale bars, 10 μm. **f**, Measurement of spontaneous excitatory postsynaptic currents (sEPSCs) in MCP-Cry2 (control)-, optoG4mt-, and optoG4-expressing mouse cultured neurons with BL stimulation. Cum., Cumulative. **g**, Schematic of viral constructs for *in vivo* optoG4 modeling and experimental schedules. **h**, Representative images of tyrosine hydroxylase (TH) (blue), αSyn (green), MCP-Cry2 (red), and *G4*-*MS2* RNA (magenta) in the nigral dopaminergic neurons exposed to BL stimulation or darkness. Scale bars, 5 μm. **i,** Representative images (left) and quantification (right) of nigral TH^+^ cells (blue) and pS129 intensity (green) in the optoG4 mouse exposed to BL stimulation or darkness. Low and high magnification scale bars, 200 and 20 μm, respectively. **j**, Motor function of optoG4 mice exposed to BL stimulation or darkness in rotarod (left) and beam-walking (right) tasks. Data are presented as mean ± standard error of mean. \**P* < 0.05 and \*\**P* < 0.01 by one-way (**f** sEPSC amplitude and frequency, and two-way (**f** (cumulative probability of sEPSC amplitude and inter-event interval), and **j**) analysis of variance with Bonferroni’s multiple comparisons test and two-sided, unpaired Student’s *t*-test (**i**). Number of replicates is shown in Supplementary Table 9.

In mouse cultured cortical neurons expressing the optoG4 system, endogenous αSyn formed pS129- and p62-positive aggregates following 3-h BL stimulation (**Fig. 4d** and **Extended Data Fig. 8g**), but did not in those expressing optoG4mt (**Fig. 4e**) or MCP-Cry2 alone (**Extended Data Fig. 8h**). Endogenous αSyn aggregation induced by optoG4 were retained even after 3-h BL stimulation and subsequent 3-h withdrawal (**Extended Data Fig. 8i**). We measured spontaneous excitatory postsynaptic currents (sEPSCs) to investigate the effect of optoG4-induced αSyn aggregation on neuronal activity. The sEPSC amplitude and frequency were significantly reduced in BL-stimulated optoG4 neurons compared to non-stimulated optoG4 neurons (**Fig. 4f**). Changes in the neuronal activities were not observed in the neurons expressing MCP-Cry2 alone or with optoG4mt, regardless of the BL conditions (**Fig. 4f** and **Extended Data Fig. 8j**). Impaired sEPSCs in the BL-stimulated optoG4 neurons were also sustained after 3-h BL withdrawal (**Extended Data Fig. 8k**).

Thereafter, we employed the optoG4 system in a series of *in vivo* experiments, injecting recombinant adeno-associated viruses (AAVs) into the nigral DA neurons of mice resulting in expression of improved Cre under the control of mouse TH promoter (AAV-mTH-iCre^42^), iCre recombinase-dependent expression of MCP-Cry2 flanked by double-floxed inverse open reading frame (DIO) (AAV-MCP-Cry2), and *G4tr*-*MS2* RNA (AAV-*G4*-*MS2*). Fourteen days after AAV injection, mice were implanted with a wireless optogenetic device to enable BL stimulation in the substantia nigra. Two days after the implantation, the mice were subjected to BL stimulation for 1-h per day for four weeks (**Fig. 4g**).

Immunohistochemical analysis confirmed MCP-Cry2 expression in the nigral DA neurons (**Extended Data Fig. 9a**). In optoG4 mice, BL stimulation formed αSyn granules, which co-assembled with MCP-Cry2 and *G4*-*MS2* RNA (**Fig. 4h**). In optoG4 mice with BL, the number of TH-positive DA neurons was significantly reduced compared to that without BL, and pS129-positive inclusions were observed (**Fig. 4i**). No pathological changes were observed in the other groups (**Extended Data Fig. 9b, c**). Rotarod and beam-walking tests were performed to determine whether optoG4 system induces PD-like motor deficits. OptoG4 mice with BL demonstrated a progressive and significant decrease in fall latency in the rotarod test and increase in the number of footslips in the beam-walking test compared to those without BL (**Fig. 4j**). The other groups did not show motor deficits, regardless of BL stimulation (**Extended Data Fig. 9d**).

### Treatment with 5-ALA ameliorates motor dysfunctions observed in PFF-injected mice

Since PPIX binds to rG4s^9^, we examined whether PPIX affects rG4s assembly *in vitro*. Treatment with PPIX reduced the size of G4tr-containing liquid droplets (**Fig. 5a**). Importantly, the formation of gel-like aggregates consisting of G4tr and αSyn was also inhibited by PPIX (**Fig. 5b** and **Extended Data Fig. 10a**).

**Fig. 5:**
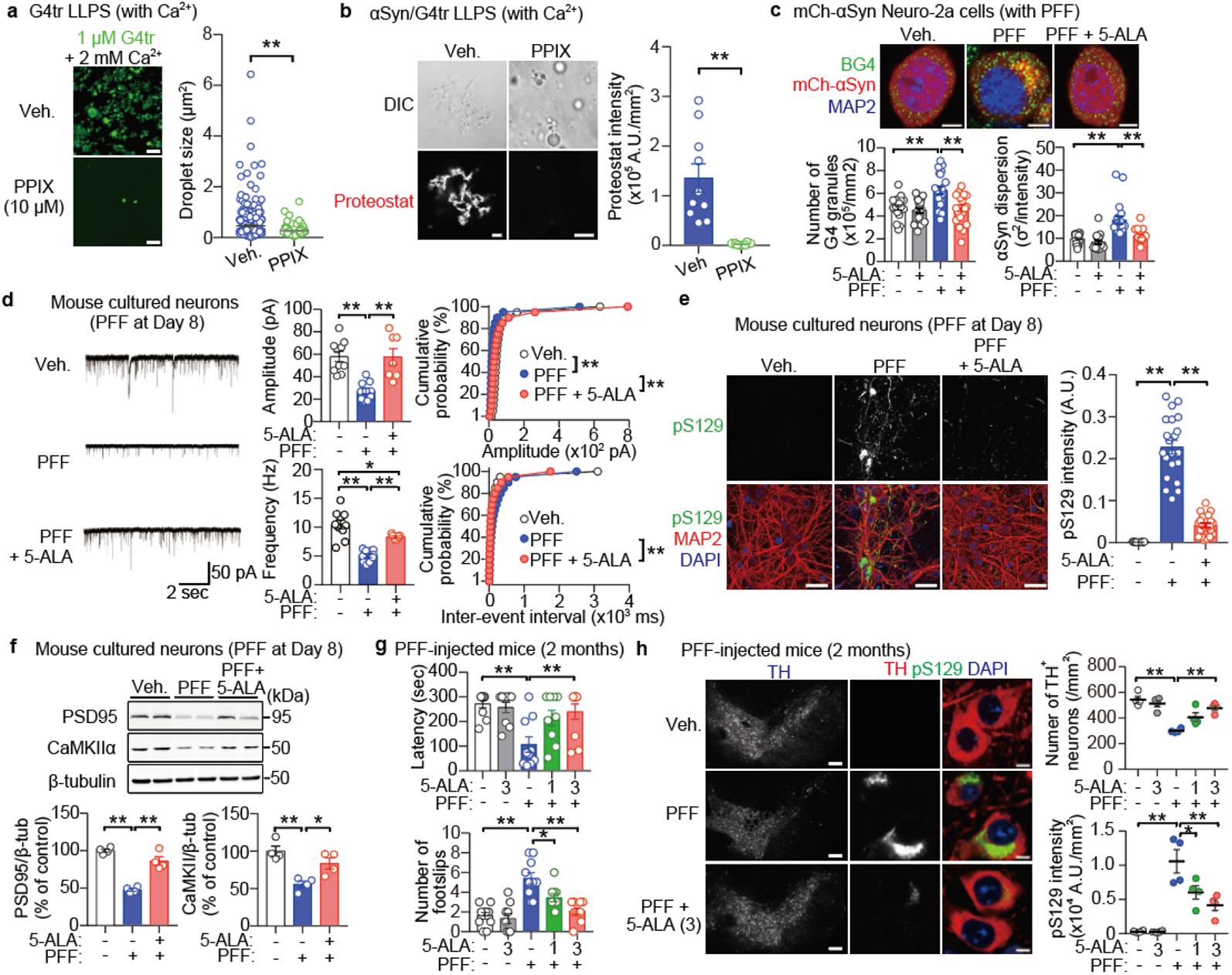
G4 ligand ameliorates neuronal dysfunction by preventing *α*-synuclein (*α*Syn) aggregation. **a**, Representative images (left) and quantification of droplet size (right) of *in vitro* phase-separated G4tr dependent on Ca^2+^ and the effect of protoporphyrin IX (PPIX). Scale bar, 5 μm. **b**, Proteostat intensity within *in vitro* phase-transitioned G4tr/αSyn dependent on Ca^2+^ and the effect of PPIX. Scale bar, 5 μm. **c**, Representative images (top) and quantification (bottom) of the number of G4 granules (green) and the dispersion of αSyn (red) in PFF-treated mCh-αSyn-expressing Neuro-2a cells and the effects of 3μM 5-aminolevulinic acid (5-ALA). Scale bars, 5 μm. **d**, Measurements of the spontaneous excitatory postsynaptic currents (sEPSCs) in PFF-treated mouse cultured neurons and the effects of 3 μM 5-ALA. **e**, Representative images (left) and quantification (right) of pS129 immunoreactivity (green) relative to MAP2^+^ area (red) in pre-formed fibrils (PFF)-treated mouse cultured neurons and the effect of 3 μM 5-ALA. Scale bars, 20 μm. **f**, Representative images (top) and quantification (bottom) of immunoblots probed with the indicated antibodies in PFF-treated mouse cultured neurons and the effects of 5-ALA. **g**, Motor function of PFF-injected mice and the effects of 5-ALA in the rotarod (top) and beam-walking (bottom) tasks. **h**, Representative images (left) and quantification (right) of the number of nigral tyrosine hydroxylase (TH^+^) cells (red) and pS129 intensity (green) relative to the TH^+^ area of PFF-injected mice and the effects of 5-ALA. Low and high magnification scale bars, 200 and 10 μm, respectively. Data are presented as mean ± standard error of mean. \**P* < 0.05 and \*\**P* < 0.01 by two-way (**d** (cumulative probability of sEPSC amplitude and inter-event interval)) and one-way (**c**,**d**,**e** (sEPSC amplitude and frequency),**f**,**g**,**h**) analysis of variance with Bonferroni’s multiple comparisons test and by unpaired two-sided, unpaired Student’s *t*-test (**a**,**b**). Number of replicates is shown in Supplementary Table 9.

PPIX is not available *in vivo* owing to its cytotoxicity^43^; thus, we investigated whether 5-ALA, an extremely low-toxic and blood–brain barrier-permeable prodrug of PPIX^9, 29^, could inhibit αSyn aggregation in cells. The application of 5-ALA significantly prevented increase in G4 granules and αSyn aggregation 24-h following PFF treatment in Neuro-2a cells expressing mCh-αSyn (**Fig. 5c** and **Extended Data Fig. 10b**). Thereafter, 5-ALA was administered to primary mouse cultured PFF-treated neurons. 5-ALA significantly restored the reduction in amplitude and frequency in sEPSC eight days after co-treatment with PFF (**Fig. 5d**). 5-ALA also reduced the pS129-positive inclusions (**Fig. 5e**), and attenuated reduction of PSD95 and CaMKIIα protein levels without changes in the mRNA levels of *Dlg4* and *Camk2a* in PFF-treated neurons (**Fig. 5f and Extended Data Fig. 10c**).

Finally, we investigated the effects of 5-ALA on αSyn aggregation and Parkinsonism *in vivo*. Mice were injected with PFF into the dorsal striatum followed by daily oral administration of 5-ALA (1 or 3 mg/kg/day). Two months after PFF injection, 5-ALA (3 mg/kg) prevented motor dysfunctions in PFF-injected mice (**Fig. 5g**). Importantly, 5-ALA also prevented the formation of pS129-positive inclusion and the resultant degeneration of nigral DA neurons (**Fig. 5h**).

## Discussion

In this study, we demonstrated that Ca^2+^-induced rG4 assembly via LLPS served as scaffolds for αSyn aggregation, contributing to neurodegeneration. rG4s bind directly to the N-terminus of αSyn, accelerating the aggregation. In PFF-treated neurons, αSyn aggregated with rG4 assembly composed of synaptic mRNAs, such as *Camk2a* and *Dlg4,* due to Ca^2+^ excess influx, eliciting neuronal dysfunction. In addition, rG4 assembly by the optoG4 system caused endogenous αSyn aggregation and neuronal dysfunction, reproducing the pathology of synucleinopathies. 5-ALA and PPIX administration suppressed rG4 LLPS and inhibited the co-aggregation of rG4s and αSyn, thereby ameliorating neurodegeneration and motor deficits in PFF-injected synucleinopathy mice. Considered together, rG4 assembly attributed to aberrant intracellular Ca^2+^ influx promotes αSyn aggregation, which may be involved in the pathogenesis of synucleinopathies **(Extended data Fig. 10).**

Although the aggregation of many neurodegenerative disease-related IDPs has been attributed to LLPS-mediated sol–gel phase transitions^22–25, 44^, purified αSyn does not form liquid droplets nor aggregates *in vitro*, except under non-physiological conditions, such as high protein concentrations above 100 μM, addition of molecular crowding reagents, low pH, and agitation (**Fig. 1**)^21, 45–47^. The difficulty of αSyn aggregation *in vitro* is presumably attributed to the monomeric state preserved by biased electrostatic and hydrophobic intramolecular interactions and intermolecular repulsion. αSyn has opposite charges at the N- and C-termini and has overall low hydrophobicity; the N-terminus with positively charged, the NAC region with high hydrophobicity, and the C-terminus with highly negatively charged that binds to bivalent metal ions, including Ca^2+48, 49^. The long-range electrostatic interactions through the N- and C-termini, as well as hydrophobic interactions between the C-terminus and NAC regions form a compact, autoinhibitory conformation that limits the exposure of the NAC region. In addition, intermolecular interactions of the negatively charged C-terminus produce electrostatic repulsion, inhibiting multimer formation^48, 49^.

Distinct from “αSyn LLPS”, we here proposed the “rG4 LLPS” hypothesis that Ca^2+^-induced association of rG4s via LLPS is responsible for the initiation of αSyn aggregation. The rG4-induced αSyn aggregation occurred under near physiological conditions (neutral pH, concentration at 34.5 μM) without both agitation and the presence of the molecular crowding reagent (**Extended Data Fig. 5**). The preferential binding of the N-terminus of αSyn to rG4s was required for αSyn aggregation **(Extended Data Fig. 4e, f**). Since the N-terminus of αSyn is involved in membrane anchoring^31^, a critical period could exist at which the interaction factor with αSyn shifts from the membrane to rG4s in the pathogenic process. Importantly, both rG4s and Ca^2+^ are necessary and sufficient for αSyn aggregation. Since rG4 unfolding mutant G4mt did not bind to αSyn, the binding of rG4s to αSyn was not through the electrostatic interaction of the negatively charged phosphate backbone of the RNAs with the positively charged N-terminus of the protein. Although how rG4 modulates αSyn conformation and the subsequent aggregation is unclear, it is possible that rG4s and Ca^2+^ shield both the N-terminus and negatively charged C-terminus, respectively, thereby exposing the NAC region upon release of the compact conformation. This would result in αSyn bearing aggregation-prone states via the hydrophobic core domain (NAC-NAC interaction). More structural analyses are warranted in the rG4s binding to the N-terminus through non-covalent bonds, such as hydrogen bonds and hydrophobic interactions using cryogenic electron microscopy and X-ray crystallography techniques.

In mammals, rG4-forming potential sequences are abundant in transcripts, with 3,800 predicted in approximately 2,300 different genes^50^. Although the rG4 assembly organizes the formation of SGs in neurons^18^, the intracellular mechanism of the rG4 assembly attributed to stress stimulation remains unclear. We here proposed that persistent and excessive intracellular Ca^2+^ influx by PFF treatment triggers rG4 assembly (**Fig. 2**). Interestingly, mRNAs encoding synaptic proteins accounted for most of the RNAs identified by BG4 RIP-seq in the PFF-treated neurons (**Fig. 3**). Consistent with this result, a significant reduction in the synaptic activity was observed in neurons challenged with optoG4 system and PFF (**Figs. 4 and 5**). Indeed, excessive Ca^2+^ influx has been identified in the pathogenesis of various neurodegenerative diseases, including synucleinopathies, Alzheimer’s disease, amyotrophic lateral sclerosis, Huntington’s disease, and spinocerebellar ataxias^32^. Furthermore, rG4-induced sol–gel phase transition of tau has been observed *in vitr*o (under preparation). These results suggest that rG4 is a key factor in the aggregation of prionoid proteins, including αSyn, tau, and FMRpolyG^9^, and excessive Ca^2+^ influx-induced rG4 assembly may be a main source of the pathogenesis in neurodegenerative diseases. In the future, it is necessary to elucidate why neuronal rG4 assembly contains a majority of synaptic mRNAs and how rG4 assembly aggregation of prionoid proteins contributes to the pathogenesis of neurodegeneration.

In conclusion, rG4 assembly by excessive Ca^2+^ influx could induce αSyn aggregation and neuronal dysfunction. In addition, we found that PPIX can inhibit the rG4 assembly, and its prodrug 5-ALA significantly ameliorated motor deficits in PFF-injected mice. Importantly, 5-ALA can act on the brain through oral administration and produce PPIX intracellularly without serious adverse effects^29^. 5-ALA can be prophylactically administered before progressive motor dysfunction, and could be a promising novel agent for neurodegenerative diseases, including synucleinopathies.

## Methods

### Animals

C57BL/6J mice (Japan SLC) were used for all the experiments. Mice were housed under climate-controlled conditions in a 12-hour light/12-hour dark cycle and were provided standard food and water *ad libitum*. Animal studies were conducted in accordance with the Kumamoto University institutional guidelines. Ethical approval was obtained from the Institutional Animal Care and Use Committee of the Kumamoto University Environmental and Safety Committee.

### Cell culture

Neuro-2A and HEK293T cell lines were grown in Dulbecco’s modified essential medium (Sigma-Aldrich) containing 10% fetal bovine serum (FBS; Gibco), 100 units/mL penicillin, and 100 μg/mL streptomycin at 5% CO_2_ and 37°C. Transfection was performed using the Lipofectamine 3,000 Reagent (Invitrogen) according to the manufacturer’s protocol. Primary cultures of mouse neurons were performed as previously described^9^. Briefly, cortical brain tissue was dissected from 18 days old embryonic mice, trypsin-digested, and mechanically dispersed. Cells were seeded with minimum essential medium (Gibco) supplemented with 10% FBS, 0.6% glucose (FUJIFILM), and 1 mM pyruvate (Sigma-Aldrich). Cells were cultured in Neuron Culture Medium (FUJIFILM) with medium changed every three days. Transfection was performed by electroporation (NEPAGENE) or Magnetofection (OZ Biosciences) according to the manufacturer’s protocol.

### Plasmid constructs

For protein purification, αSyn constructs with mouse coding sequence (CDS), mutations, and CDS of fluorescent proteins were subcloned into pET-human αSyn^51^ using the KOD-Plus Mutagenesis kit (TOYOBO) and In-Fusion HD Cloning Kit (Takara Bio) according to the manufacturers’ protocol. For preparing the mammalian expression plasmids, pCAG-GFP^9^, pHR-FUSN-mCh-Cry2WT (Addgene #101223), and MCP-YFP (Addgene #101160) were used as backbones. Repeat sequences of (TTAGGG)_23_ and (TTACCG)_23_ were synthesized by GeneScript and subcloned into pHR-Tre3G-29×GGGGCC-12×MS2 (Addgene #99149) using Ligation high (TOYOBO). The fragments with MS2 were inserted in the 3’-untranslated region of pCAG-HaloTag vector^9^. For recombinant AAV production, pAAV-TH-iCre, pAAV-hSyn-DIO-MCP-Cry2, pAAV-hSyn-G4tr-MS2, and pAAV-hSyn-G4mt-MS2 were synthesized by GeneScript, using pAAV-hSyn-DIO{ChETA-mRuby2}on-W3SL (Addgene #111389) as a backbone.

### Antibodies

The primary and secondary antibodies for immunoblotting and immunohistochemical analyses utilized in the present study are shown in the Supplementary Method.

### Preparation of recombinant proteins

pET-αSyn plasmids were transformed into *Escherichia coli* BLR (DE3) cells and purified as described previously^52^. Briefly, bacterial lysate by ultrasonication was heated (75°C, 15 min) and precipitated by ammonium sulfate. The dialysate was fractionated by ion exchange chromatography (Resource-Q or HiTrap SP HP; GE Healthcare) on an AKTA-explorer system (GE Healthcare) at 4°C. Samples were desalted, lyophilized, and stored at 4°C until the assay was conducted.

### Preparation of *α*Syn PFF

αSyn PFF was prepared as previously described^51^. Briefly, recombinant αSyn (5 mg/mL in 1× phosphate buffer saline (PBS)) was centrifuged for an hour at 100,000g at 4°C and the supernatants were shaken in an orbital thermomixer at 1,000 rpm for seven days. αSyn PFFs were sonicated by Bioruptor (Cosmo Bio) for eight cycles at high power (on, 1-pulse every other second for 15 s in total; off, 15 s; at 23°C), and then stored at – 80°C until use.

### AAV preparation

Recombinant AAV9 vectors were generated by co-transfection of AAVpro 293T cells (Takara Bio) with three plasmids: pAAV, pHelper (Stratagene), and pAAV2/9 (a gift from James M. Wilson). The viral particles were purified from transfected cells and titrated using the AAVpro Purification Kit Maxi and AAVpro Titration Kit (Takara Bio) according to the manufacturer’s instructions.

### Treatment with *α*Syn PFF and other compounds in cultured cells

Neuro-2a or HEK293T cells expressing mCh- or GFP-human αSyn were treated with human αSyn PFF at 50 μg/mL (3.45 μM) for the indicated time. Mouse cultured neurons at DIV10 were incubated with mouse or human αSyn PFF at 50 μg/mL for the indicated time, some of which were further treated with 5 μM Calbryte 520AM (AAT Bioquest) for 30 min. Several mouse cultured neurons were treated with 0.5 mM NaAsO_2_ (Sigma-Aldrich) for 60 min or 1.0 μM ionomycin (AdipoGen Life Sciences) for 30 min on DIV14.

### Intracellular Ca^2+^ imaging

αSyn PFF- and Calbryte 520AM-treated mouse cultured neurons were subjected to measurement of intracellular Ca^2+^ levels using LSM900 microscopy system (Carl Zeiss) in an imaging chamber (KYODO INTERNATIONAL).

### Stereotaxic injection of *α*Syn PFF and AAV

Mouse αSyn PFFs were injected into the bilateral dorsal striatum (2 μg/hemisphere) at the following coordinates (in mm): anterior, +0.7; lateral, ±2.5; and depth, −2.8 relative to the bregma. Viral particles of 1-μL with the same titer (1.4 × 10^10^ vector genomes/mL) were injected into the bilateral substantia nigra at the following coordinates (in mm): anterior, −3.5; lateral, ±1.2; and depth, −4.0 relative to the bregma. αSyn PFFs and AAV were injected at 0.2 µL/min using a Hamilton syringe under 1% isoflurane anesthesia.

### FRAP assay in living cells

GFP-αSyn-transfected and αSyn PFF-transduced HEK293T cells were subjected to FRAP assay in the imaging chamber. Fluorescent images were obtained using the LSM900 microscopy system. Photobleaching was performed with 50% laser power with 20 iterations. Time-lapse images were recorded every 250 ms at room temperature.

### *In vitro* LLPS, Proteostat, and FRAP assay

Non-labeled or mCh-labeled human αSyn (34.5 or 69 μM) was prepared in the LLPS assay buffer: 50 mM Tris-HCl buffer (pH 7.5) containing 140 mM KCl, 15 mM NaCl, 10 mM MgCl_2_, 0–20% PEG8000 (MP Biomedicals), and 1 unit/μL RNase inhibitor (TOYOBO) in the presence or absence of 2 mM CaCl_2_ and 2.5 mM EGTA. Total protein, total RNA, or synthesized 24-mer RNA oligomer (1 μM; Hokkaido System Science) was added and incubated at 37℃ for three hours. Total protein was lysed by homogenization in radioimmunoprecipitation (RIPA) buffer (FUJIFILM) containing protease inhibitor cocktail (NACALAI TESQUE) and Cryonase Cold-active Nuclease (TAKARA), while total RNA was extracted using an RNeasy Mini Kit (Qiagen), both from intact Neuro-2a cells. The resultant solutions were mounted on glass slides using a 0.12 nm spacer (Sigma-Aldrich) and a coverslip. In some conditions, 2% (v/v) Proteostat reagent (Enzo Life Sciences) was added into the LLPS assay buffer to detect aggregates. Differential interference contrast images were obtained using the TCS SP8 confocal microscopy system (Leica Microsystems). Samples were photobleached with 50% laser power and time-lapse images were recorded every 8-s using the LSM780 microscopy system.

### Circular dichroism (CD) spectroscopy

G4tr (UUAGGG)_4_ and G4mt (UUACCG)_4_ were prepared using an *in vitro* the HiScribe T7 Quick High Yield RNA Synthesis Kit (New England Biolabs) following the manufacturers’ instructions, respectively. Other RNA oligomers were synthesized by Hokkaido System Science. Each RNA (2.5 μM) was prepared in 10 mM Tris-HCl buffer (pH 7.5) containing 100 mM LiCl or KCl. The oligomers were then refolded by a heating/cooling process (90°C for 3 min, followed by cooling down to 10°C for 1.5 hours) before measurement. The CD spectra were recorded at room temperature over the range of 200–350 nm using a spectrometer (J-805LST; JASCO) equipped with a 3 mm path-length quartz cuvette.

### EMSA

Fluorescein phosphoramidite (FAM)-labeled RNAs (20 nM) in 10 mM Tris-HCl buffer (pH 7.5) containing 100 mM NaCl were refolded as described above. The folded RNA samples were incubated with a human αSyn (0–138.4 μM) or BG4 (320 nM) with 1 unit/μL RNase inhibitor (TOYOBO) at room temperature for at least 30 min. The resulting mixtures were analyzed by 6% native TBE polyacrylamide gel electrophoresis and visualized using Typhoon Trio equipment (GE Healthcare).

### SPR-binding assays

SPR experiments were performed on OpenSPR (Nicoya). After biotin-labeled G4tr or G4mt was captured by a sensor chip using biotin-streptavidin sensor kit (Nicoya), αSyn in running buffer (50 mM Tris-HCl pH 7.5 and 10 mM KCl) was streamed over the sensor chip to allow interaction. The interaction time was at least 5 min to allow the ligand signal to stabilize. The running buffer was then flowed at a rate of 20 μL/min for 5 min to collect the dissociation data. Binding kinetic parameters were obtained by fitting the curve to a One-to-One binding model using TraceDrawer software (Ridgeview Instruments).

### RNA Bind-n-Seq

RNA Bind-n-Seq was performed as previously described^30^. Briefly, the purified αSyn (0–69 μM) and BG4 (0.32 μM) were equilibrated in binding buffer (25 mM Tris-HCl pH 7.5, 150 mM KCl, 3 mM MgCl_2_, 0.01% Tween 20, 1 mg/ml bovine serum albumin (BSA), and 1 mM dithiothreitol) with 1 units/μL RNase inhibitor (TOYOBO) for 30 min at room temperature. 24-mer random RNAs of 1 μM (Hokkaido System Science) were added to the solutions and incubated for an hour at 4℃. During the incubation, Dynabeads Protein G (Invitrogen) was washed with wash buffer (25 mM Tris-HCl pH 7.5, 150 mM KCl, 60 μg/mL BSA, 0.5 mM EDTA, and 0.01% Tween 20) and then equilibrated in binding buffer including 1 units/μL RNase inhibitor with anti-αSyn or anti-DYKDDDDK tag antibody until use. The protein/RNA complex was pulled down by the beads and incubated for an hour at room temperature. Supernatant was removed and the beads were washed once. The beads were incubated at 70℃ for 10 min in elution buffer (10 mM Tris-HCl pH 7.0, 1% sodium dodecyl sulfate (SDS), and 1 mM EDTA). Bound RNA was extracted and reverse-transcribed into cDNA using Superscript III (Invitrogen) according to the manufacturer’s instructions together with the following primer: 5’-GCCTTGGCACCCGAGAATTCCA-3’. To control for any nucleotide biases in the input random library, the input RNA pool (0.5 pmol) was also reverse-transcribed and Illumina sequencing library prep was followed by 8–10 cycles of polymerase chain reaction (PCR) using High Fidelity Phusion (New England Biolabs). All the libraries were barcoded in the PCR step, pooled together, and sequenced on a NextSeq 500 sequencer. Motif enrichment analyses were performed by the MERMADE program^53^. Base contents, GC contents, and GC skew were calculated by the SeqKit program^54^. The motifs were processed by enoLOGOS21 (http://www.benoslab.pitt.edu/cgi-bin/enologos/enologos.cgi), and putative G4 sequences were predicted using QuadBase2 (https://quadbase.igib.res.in/).

### RIP-seq

Immunoprecipitation of RNPs was performed using a RiboCluster Profiler RIP-Assay kit for microRNA (MBL) according to the manufacturer’s protocol. Cells were harvested, lysed, and precleared on Dynabeads Protein G for an hour at 4℃. The precleared lysates were incubated at 4°C for three hours in the beads immobilized with normal rabbit IgG, anti-αSyn, or BG4 with anti-DYKDDDDK tag antibody. Each fraction containing protein-bound RNA was purified and prepared for RNA-sequencing library using NEBNext Ultra II Directional RNA Library Prep Kit for Illumina (New England Biolabs). All the libraries were sequenced on a NextSeq 500 sequencer (Illumina). Fastq files were processed for quality check and adapter trimming with the Trimgalore! (v0.6.6). Trimmed reads were used for mapping the mouse genome (mm10) using the STAR (v2.7.9a). DESeq2(v1.38.3) was used for identifying BG4-enriched transcripts. Putative rG4-forming motifs of mRNAs were predicted using the QGRS mapper (http://bioinformatics.ramapo.edu/QGRS/index.php). GO analysis was performed using DAVID Bioinformatics Resources (https://david.ncifcrf.gov/home.jsp).

### Photoactivation by exposure of BL stimulation

For live cell imaging, OptoG4-expressing Neuro-2a cells were placed in the imaging chamber and imaged typically with two laser wavelengths (488 nm for Cry2olig photoactivation/560 nm for mCh imaging) using the LSM780 microscopy system. Cry2olig photoactivation was induced at 5% power and time-lapse acquisitions were performed every 20 s at room temperature. For electrophysiological and immunohistochemical analyses, cells were photoactivated using 470 nm blue light-emitting diode (LED) light (Bio Research Center) with an estimated light intensity of 1.0 mW/cm^2^. For *in vivo* photoactivation, mice were implanted with dual-LED optic cannulae at 470 nm (Bio Research Center) into the bilateral substantia nigra two weeks following AAV injection. Daily photoactivation was performed using Teleopto, a wireless optogenetic stimulation system (Bio Research Center).

### Drugs

PPIX was purchased from Sigma-Aldrich. 5-ALA hydrochloride and sodium ferrous citrate were provided by SBI Pharmaceuticals. PPIX (10 μM) was added to the LLPS assay buffer. Cultured cells were co-treated with PFF and 5-ALA (3 μM with sodium ferrous citrate [20:1 molar ratio] dissolved in sterilized distilled water) or vehicle (sodium ferrous citrate). Animals were orally administered with vehicle or 5-ALA (1 or 3 mg/kg/day) with sodium ferrous citrate 24 hours following αSyn PFF injection daily for two months.

### Electrophysiology

sEPSC was recorded at room temperature for mouse cultured neurons on DIV14 (optoG4) or DIV18 (αSyn PFF) using an EPC10 amplifier (HEKA Instruments) as previously described^9^. The following buffers were used in the present study: extracellular buffer (143 mM NaCl, 5 mM KCl, 2 mM CaCl_2_, 1 mM MgCl_2_, and 10 mM glucose, and 10 mM HEPES at pH 7.4 adjusted with NaOH) and intracellular buffer (135 mM CsMeS, 5 mM CsCl, 0.5 mM EGTA, 1 mM MgCl_2_, 4 mM Mg_2_ATP, 0.4 mM NaGTP, 5 mM QX-314 and 10mM HEPES at pH 7.4 adjusted with CsOH). Recording pipettes had a resistance of 3.5–4.5 MΩ when filled with the intracellular buffer. The sEPSC was recorded for 2 min at a holding potential of −70 mV in the presence of 10 μM bicuculline in the extracellular buffer. Recordings were filtered at 2 kHz and digitized at 10 kHz. Data were collected and initially analyzed using Patchmaster software (HEKA). Further analysis was performed using SutterPatch v2.2 (Sutter Instrument).

### Immunohistochemical analysis

Immunohistochemistry was performed as previously described^9^. Cells and mouse brain slices (50 μm thickness) were fixed with 4% paraformaldehyde, washed with 1× PBS, permeabilized with 0.1–0.3% Triton X-100, and incubated with 1% BSA and 0.1–0.3% Triton X-100 in 1× PBS. Human frozen brain sections (5–10 μm thickness) were obtained from BioChain Institute and a part of them were treated with RNase A. An hour after blocking, samples were treated with primary antibodies for 1–3 days at 4°C. After incubation with fluorophore-conjugated secondary antibodies, samples were mounted in Vectashield (Vector Laboratories). Some samples were counterstained with 4’,6-diamidino-2-phenylindole. Immunofluorescence images were obtained using LSM900 or TCS SP8 confocal microscopy system.

### Fluorescent *in situ* hybridization (FISH)

FISH was performed as previously described^9^. After permeabilization as described above, the samples were prehybridized with hybridization solution (2× saline-sodium citrate (SSC), 40% formamide, 10% dextran sulfate, 2 mM vanadyl sulfate, 0.5 mg/mL yeast transfer RNA) for 60 min at 37 (*MS2*) or 60°C (*Camk2a* and *Dlg4*), and then incubated with 1 nM (*MS2*) or 0.3 μM (*Camk2a* and *Dlg4*) probe in a hybridization solution at 37 or 60°C overnight. After hybridization, samples were washed and subjected to immunofluorescence procedures and images were collected as described above. FISH probe sequences are shown in the Supplemental Method.

### Western blotting analysis

Western blot analyses were performed as previously described^9^. Briefly, primary mouse cultured neurons were homogenized in RIPA buffer with a protein inhibitor cocktail. The lysate was centrifuged and protein concentration of the supernatant was determined by BCA method, and samples were then boiled for 3 min in Laemmli’s sample buffer. Equal amounts of protein were loaded onto SDS-polyacrylamide gels. After transferring the separated proteins to polyvinylidene difluoride membranes, the membranes were blocked with 5% non-fat skim milk and incubated with primary antibody at 4°C overnight. Blots were developed using an ECL Western Blotting Substrate (Thermo Scientific), and the signals were detected using LAS-4000mini (GE Healthcare).

### Quantitative PCR with reverse transcription (RT-qPCR)

Sample preparation for RT-qPCR from primary cultured neurons was performed using the RNeasy Mini Kit and PrimeScript RT Master Mix (Takara Bio). RT-qPCR was performed using KOD SYBR qPCR Mix (TOYOBO) on a CFX Connect Real-Time PCR System (Bio-Rad Laboratories). Gene expression was assessed using the differences in normalized Ct (cycle threshold) (^ΔΔ^Ct) method after normalization to *Gapdh* expression. Fold change was calculated by 2^-ΔΔCt^. PCR primers are shown in the Supplemental Method.

### Behavioral analysis

The rotarod and beam-walking tasks were performed according to a previous report^51^. Briefly, mice were trained before the stereotaxic injection. In the rotarod task, mice were placed on the rod rotating at 20 rpm and falling latency was measured for up to 5 min. In the beam-walking task, the number of foot-slips from the end of beam to goal box was recorded.

### Statistical analysis

Data are presented as the mean ± standard error of the mean. Comparisons between two experimental groups were made using unpaired Student’s t-test. Statistical significance for differences among groups was tested by one-way or two-way analysis of variance with *post-hoc* Bonferroni’s multiple comparison test. *P* < 0.05 represented a statistically significant difference. All the statistical analyses were performed using GraphPad Prism 7 (GraphPad Software). Source data are provided in Supplementary Table 9.

## Acknowledgments

This work was supported by AMED (grant numbers: JP21wm0525023 and JP21gm6410021 to Y.Y.; JP22ek0109591 and JP23ek0109651 to N.S.), JSPS KAKENHI (grant numbers: JP21K20723 and JP22J00687 to K.Matsuo; JP21K06579 to Y.Y.; and JP21H00207, JP20K21400, and JP22K19297 and JP23H00373 to N.S.), JST FOREST Program (JPMJFR2043 to N.S.), The Pharmacological Research Foundation, Tokyo (to Y.Y.), Kowa Life Science Foundation (to Y.Y.), Narishige Neuroscience Research Foundation (to Y.Y.), the Astellas Foundation for Research on Metabolic Disorders (to N.S.), the Foundation of SBI Pharmaceuticals Co., Ltd. (to N.S.), International Research Core for Stem Cell-based Developmental Medicine Encourage Project and The Inter-University Research Network for High Depth Omics, Institute of Molecular Embryology and Genetics, Kumamoto University.

## Author Contributions

N.S. and Y.Y. designed the study. K.Matsuo, S.A., K.Maeda, K.Kawakubo, G.K., K.Kudo, Y.S., K.H., S.I., S.U., S.F., and Y.Y. performed the experiments. Y.K. and T.M. provided critical advice for αSyn synthesis and analysis of the interaction between rG4 and αSyn. K.Matsuo, N.S., and Y.Y. wrote the manuscript. N.S. and Y.Y. supervised the project.

## Data Availability

The raw data of RNA-Bind-n-Seq and RIP-Seq analyses are available at Gene Expression Omnibus (accession number: GSE235418 and GSE234238, respectively). Additional data related to this paper may be requested from the corresponding author N.S. and Y.Y. upon reasonable request.

Supplementary Information is available for this paper

---

**Extended Data Fig. 1:**
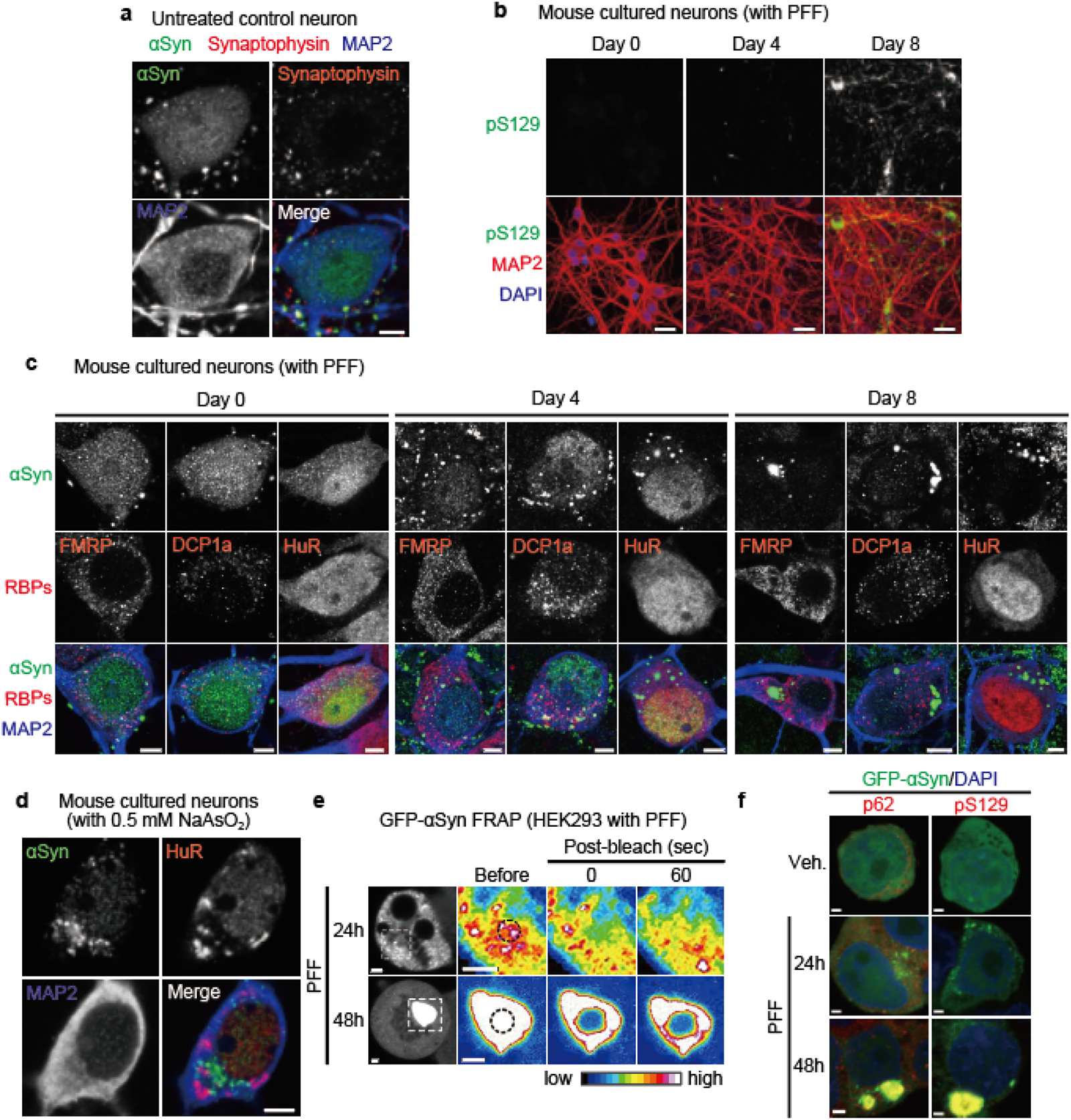
Localization and dynamics of *α*-synuclein (*α*Syn) under cellular stress. **a**, Representative images of αSyn (green), synaptophysin (red), and MAP2 (blue) in naïve mouse cultured neurons. Scale bar, 5 μm. **b**, Representative images of pS129 (green), MAP2 (red), and 4’,6-diamidino-2-phenylindole (DAPI) (blue) in mouse cultured neurons following pre-formed fibrils (PFF) treatment for the indicated time. Scale bars, 20 μm. **c**, Representative images of αSyn (green), RNA binding proteins (red), and MAP2 (blue) in mouse cultured neurons after PFF treatment for the indicated time. Scale bars, 5 μm. **d**, Representative images of αSyn (green), HuR (red), and MAP2 (blue) in mouse cultured neurons after treatment with 0.5 mM NaAsO_2_ for 60 min. Scale bar, 5 μm. **e,** Representative images of αSyn fluorescence recovery after photobleaching assay in PFF-treated GFP-αSyn expressing HEK293T cells. Scale bars, 2 μm. **(f)**, Representative images of p62 or pS129 (red) and DAPI (blue) in HEK293T cells treated with PFF following GFP-αSyn overexpression (green). Scale bars, 2 μm.

**Extended Data Fig. 2:**
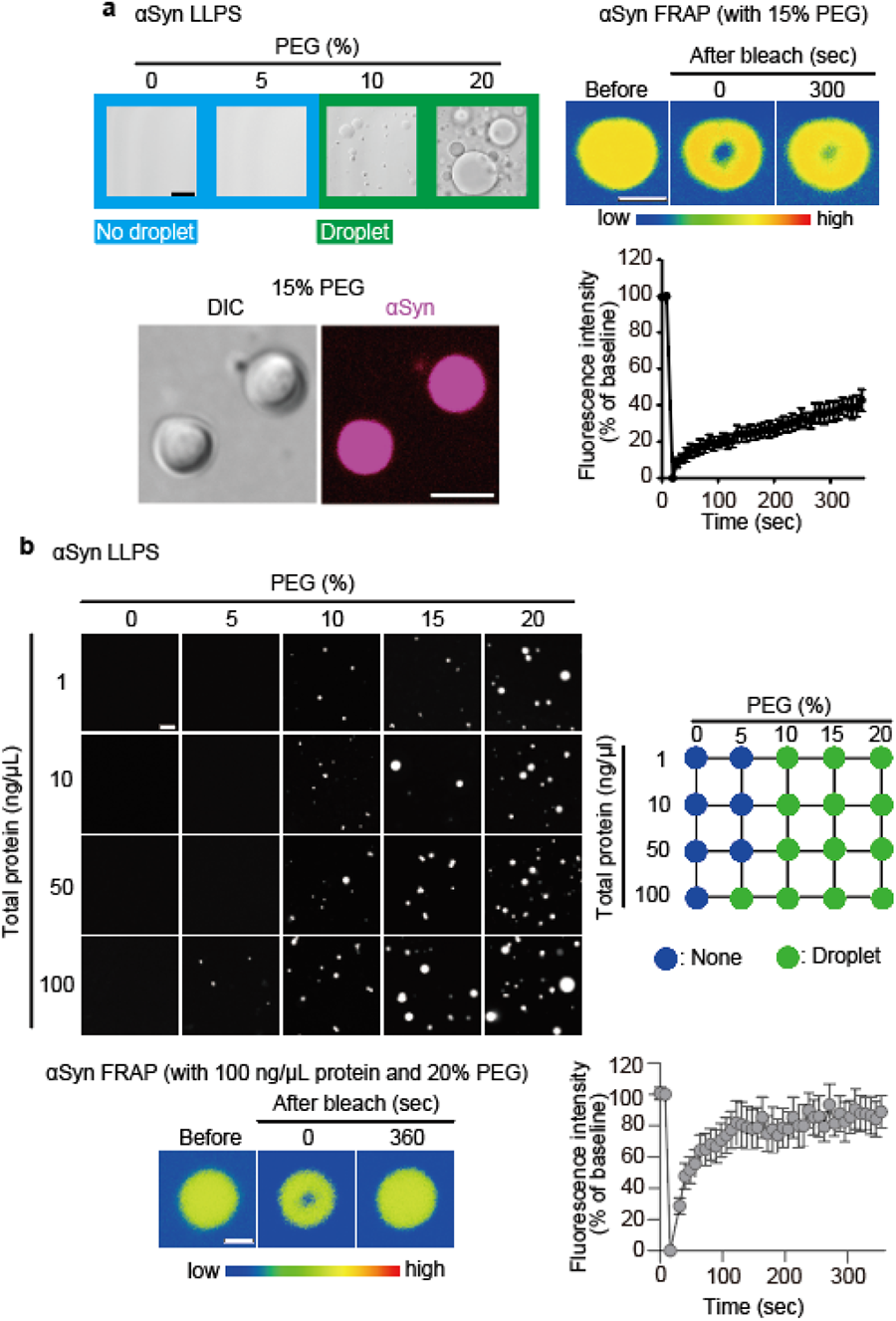
*In vitro α*-synuclein (*α*Syn) phase separation in the presence of a molecular crowding agent. **a**, *In vitro* αSyn phase separation (left) and fluorescence recovery after photobleaching (FRAP) assay (right) dependent on polyethylene glycol (PEG). Scale bars, 5 (Differential interference contrast image) and 2 (Fluorescent images) μm, respectively. **b**, *In vitro* αSyn phase separation (top) and FRAP assay (bottom) dependent on PEG and total protein extracted from Neuro-2a cells. Data are presented as mean ± standard error of mean. n = 7 (**a**) and n = 7 (**b**).

**Extended Data Fig. 3:**
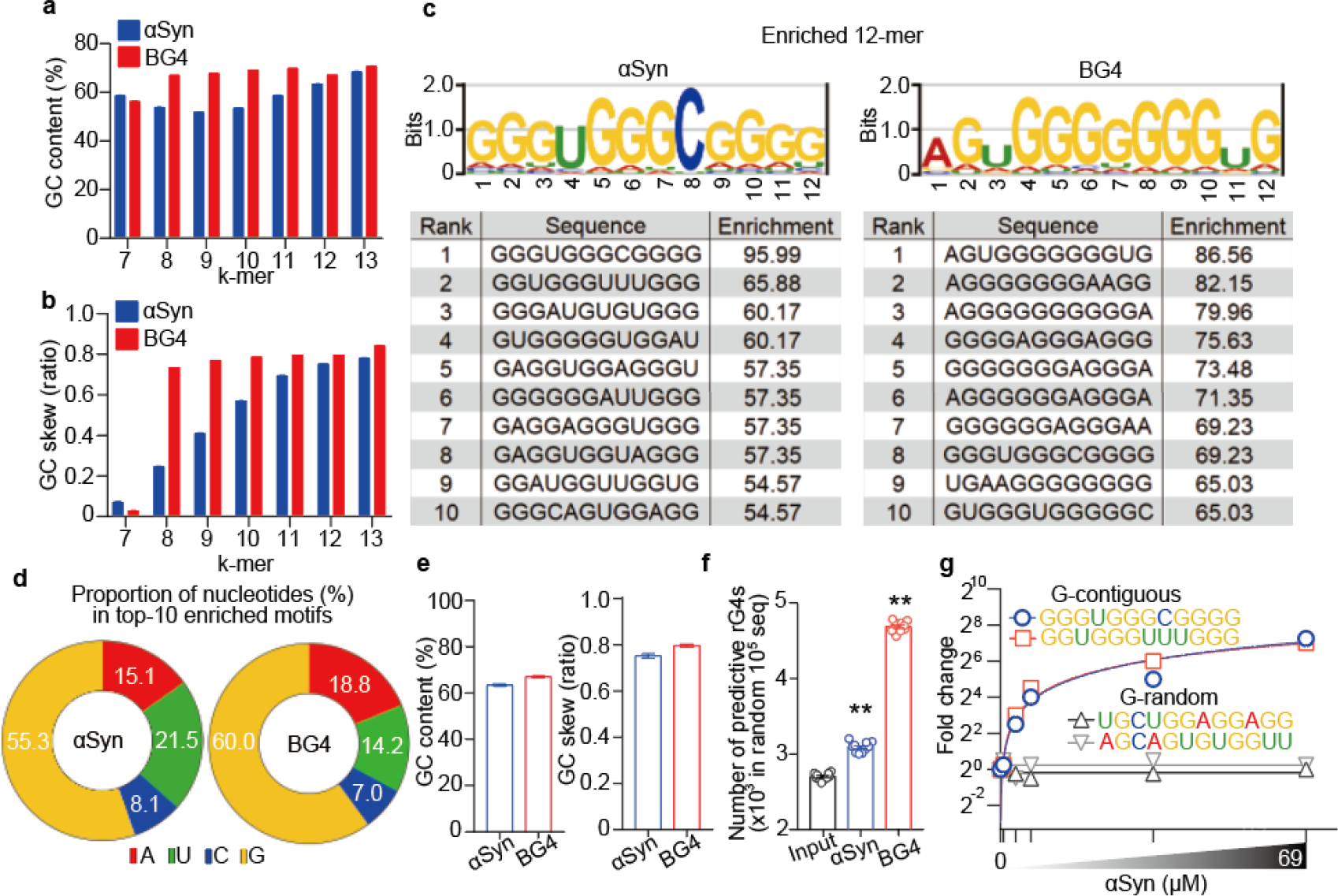
Analysis of RNA Bind-n-Seq for *α*-synuclein (*α*Syn). **a**,**b**, GC contents (**a**) and GC skew (**b**) in *k*-mer sequences in enriched motifs for αSyn and BG4. **c**–**e**, The top 10 motifs enriched in αSyn and BG4 (**c**), the proportion of nucleotides (**d**), and the GC contents and GC skew (**e**). **f**, Number of sequences with predicted rG4 structures enriched in αSyn and BG4. **g**, Fold enrichment of the top two 12-mers (circle and square marks) and two randomly chosen 12-mers (triangle marks) across αSyn concentrations (see legends in Fig. 1f). Data are presented as mean ± standard error of mean. \*\**P* < 0.01 by one-way analysis of variance with Bonferroni’s multiple comparisons test. Number of replicates is shown in Supplementary Table 9.

**Extended Data Fig. 4:**
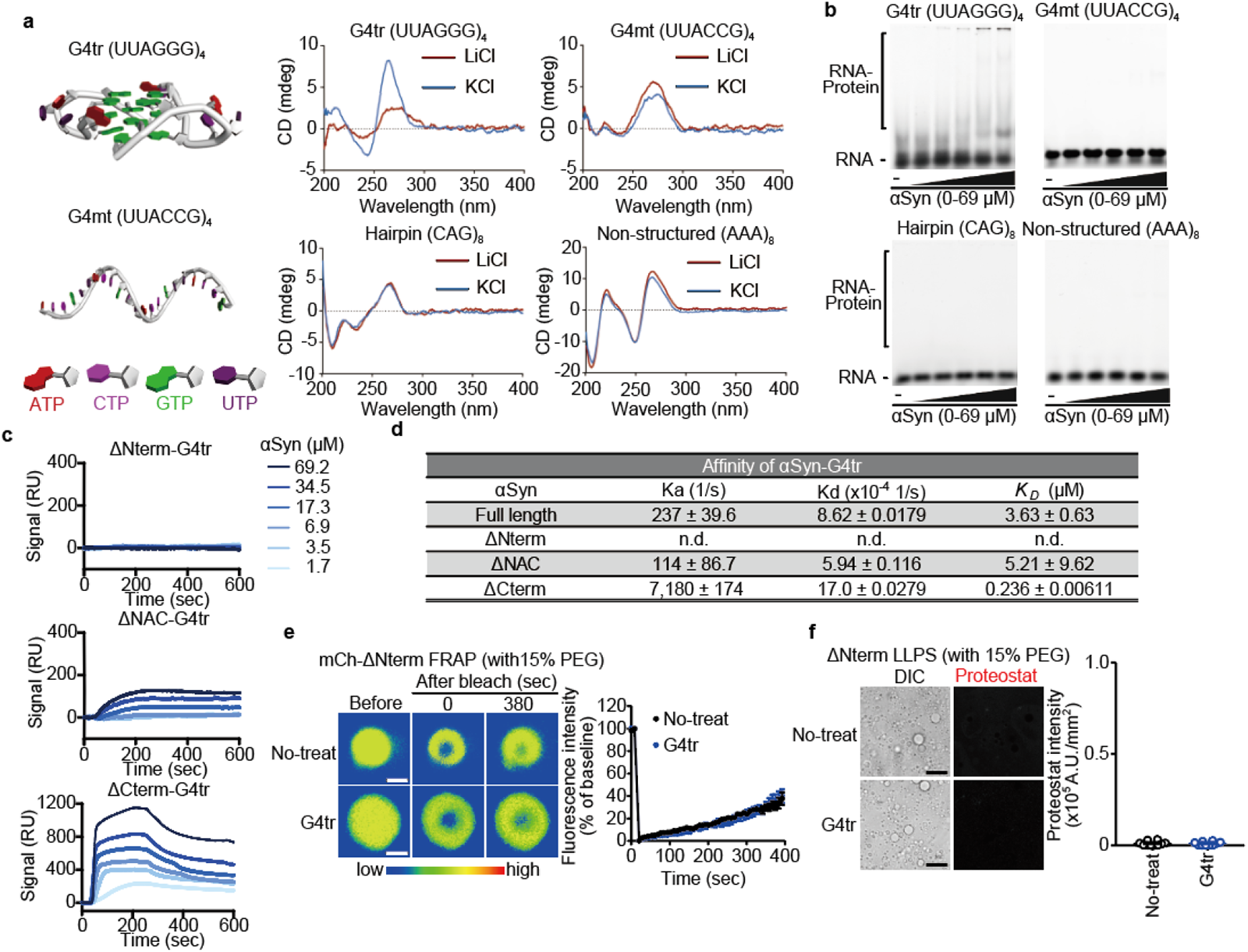
Physicochemical and biochemical analyses of RNA structures interacting to *α*-synuclein (*α*Syn). **a**, Schematic illustration of G4tr and G4mt (left). Circular dichroism spectra of the indicated RNAs in the presence of 100 mM KCl or LiCl (right). **b**, Representative images of electrophoresis mobility shift assay for the interaction of αSyn (0, 0.69, 3.45, 6.9, 34.5, and 69 μM μM) with Fluorescein phosphoramidite-labeled G4tr, G4mt, (CAG)_8_, or (AAA)_8_ oligomers (20 nM) in the presence of 100 mM NaCl. **c**,**d**, Representative surface plasmon resonance binding curves (**c**) and the detailed parameters (**d**) related to the interaction of G4tr with αSyn ΔNterm, ΔNAC, and ΔCterm. *K_D_*, dissociation constant; Ka, association rate constant; Kd, dissociation rate constant; n.d., not detected. **e**,**f**, *In vitro* G4tr/αSyn ΔNterm fluorescence recovery after photobleaching assay (**e**) and Proteostat intensity within the phase-separated droplets (**f**) dependent on 15% PEG. Scale bars, 1 (**e**) and 10 (**f**) μm, respectively. Data are presented as mean ± standard error of mean. Number of replicates is shown in Supplementary Table 9.

**Extended Data Fig. 5:**
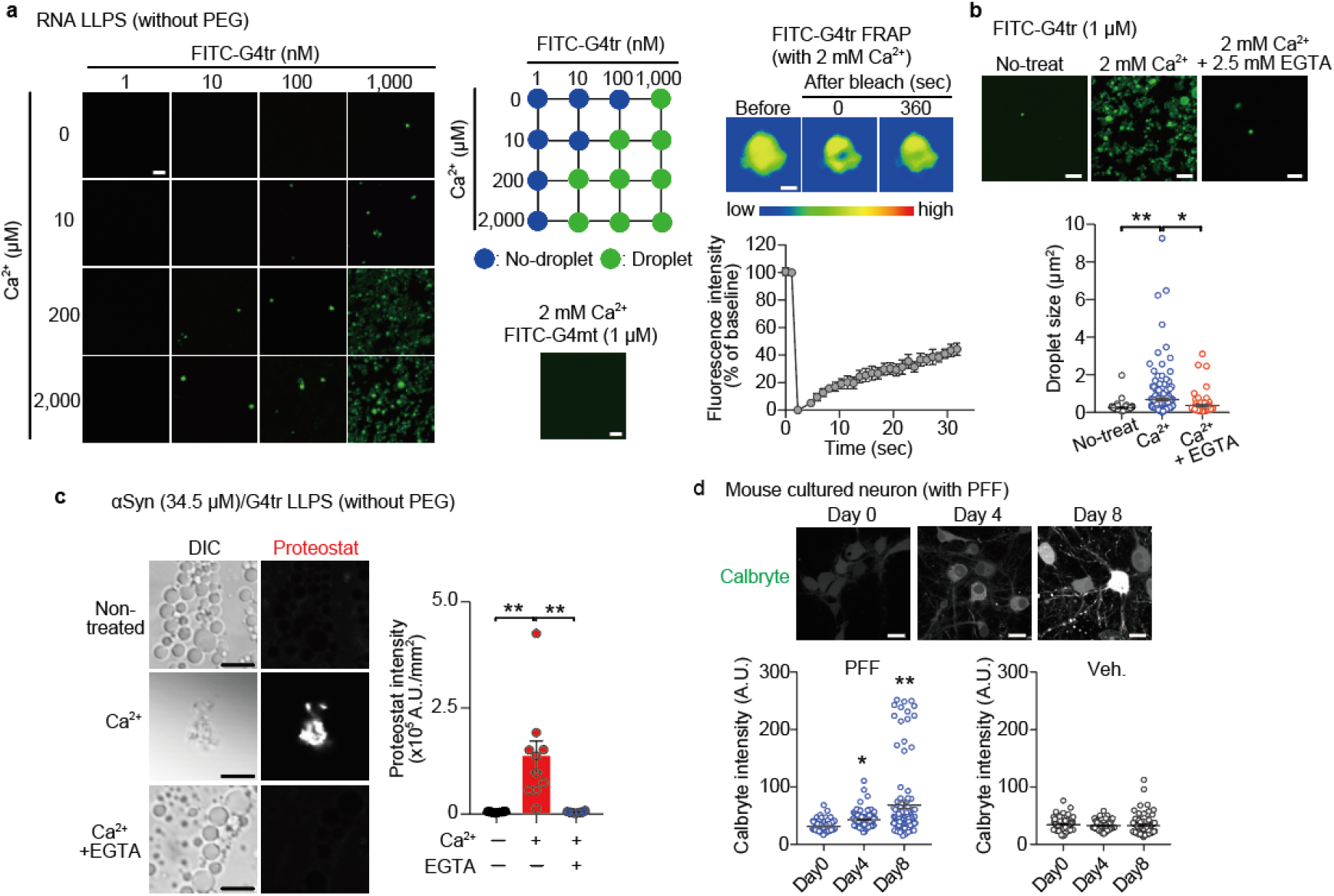
Ca^2+^-dependent G4tr assembly promotes *α*-synuclein (*α*Syn) sol–gel phase transition. **a**, *In vitro* G4tr and G4mt phase separation (left, center) and fluorescence recovery after photobleaching (FRAP) assay (right) dependent on Ca^2+^. Scale bars, 5 μm. n = 5 (FRAP). **b**, Representative images (top) and quantification of droplet size (bottom) of *in vitro* phase-separated G4tr dependent on Ca^2+^ and the effect of EGTA. Scale bars, 5 μm. **c**, Representative images (left) and quantification (right) of Proteostat intensity within phase-transitioned G4tr/αSyn dependent on Ca^2+^. Scale bars, 5 μm. **d**, Representative images (top) and quantification (bottom) of Calbryte 520AM intensity in mouse cultured neurons after PFF treatment for the indicated time. Scale bars, 20 μm. Data are mean ± s.e.m. \**P* < 0.05 and \*\**P* < 0.01 by one-way analysis of variance with Bonferroni’s multiple comparisons test. Number of replicates is shown in Supplementary Table 9.

**Extended Data Fig. 6:**
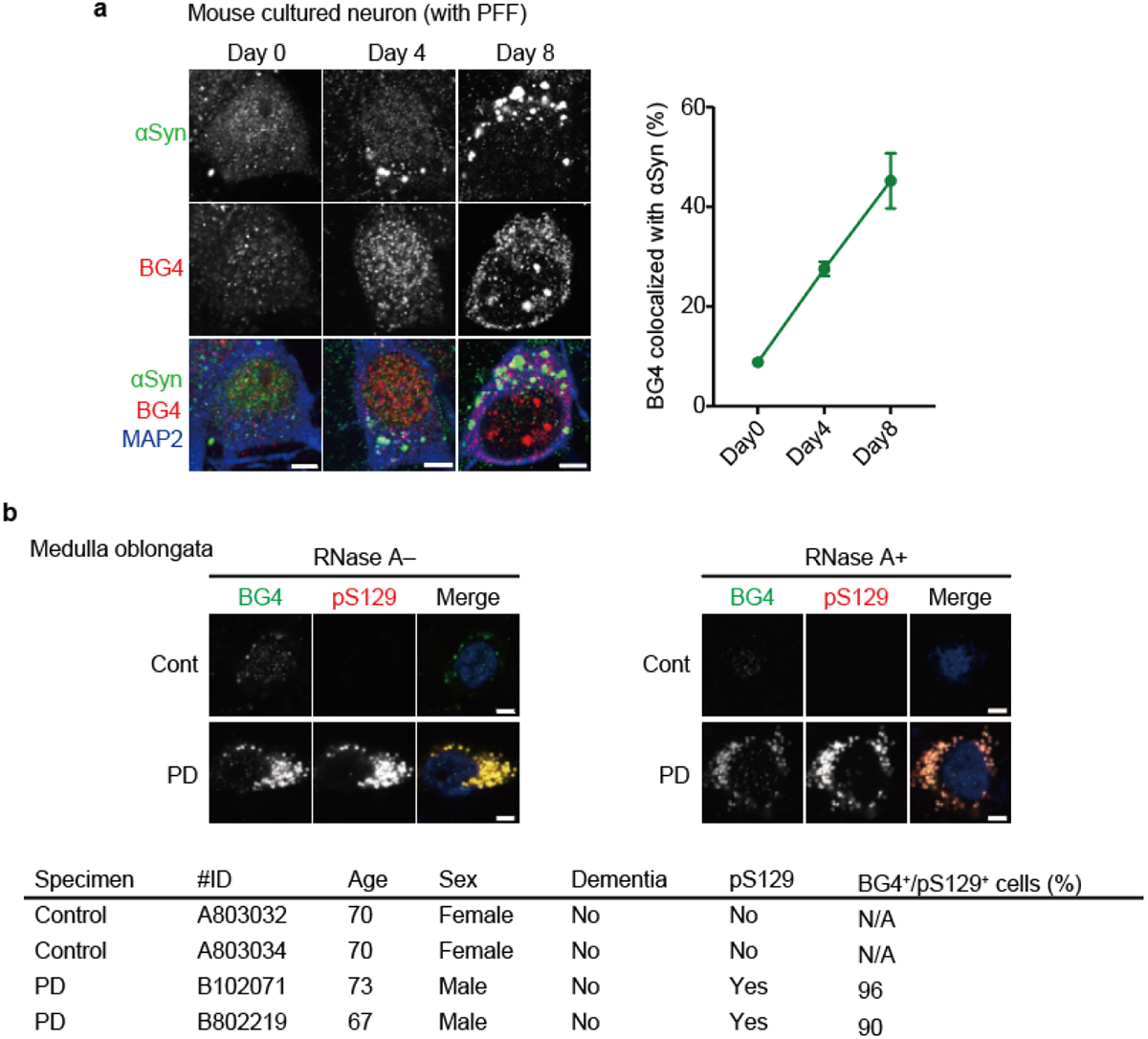
Co-aggregation of RNA G-quadruplexes and *α*-synuclein (*α*Syn) in pre-formed fibrils (PFF)-treated mouse cultured neurons and postmortem brain of patients with Parkinson’s disease. **a**, Representative images (left) and quantification (right) of co-localization of αSyn (green) and BG4 (red) granules in MAP2^+^ cells (blue) in mouse cultured neurons after PFF treatment. n = 12 (day 0), 15 (day 4), and 8 (day 8). Scale bars, 5 μm. **b**, Representative images of BG4 (green), pS129 (red), and 4’,6-diamidino-2-phenylindole (blue) in the medulla oblongata of human normal control and patients with PD treated with or without RNase A (top). Scale bars, 5 μm. Information for human samples and the proportion of BG4^+^ cells among pS129^+^ cells (bottom). Data are presented as mean ± standard error of mean.

**Extended Data Fig. 7:**
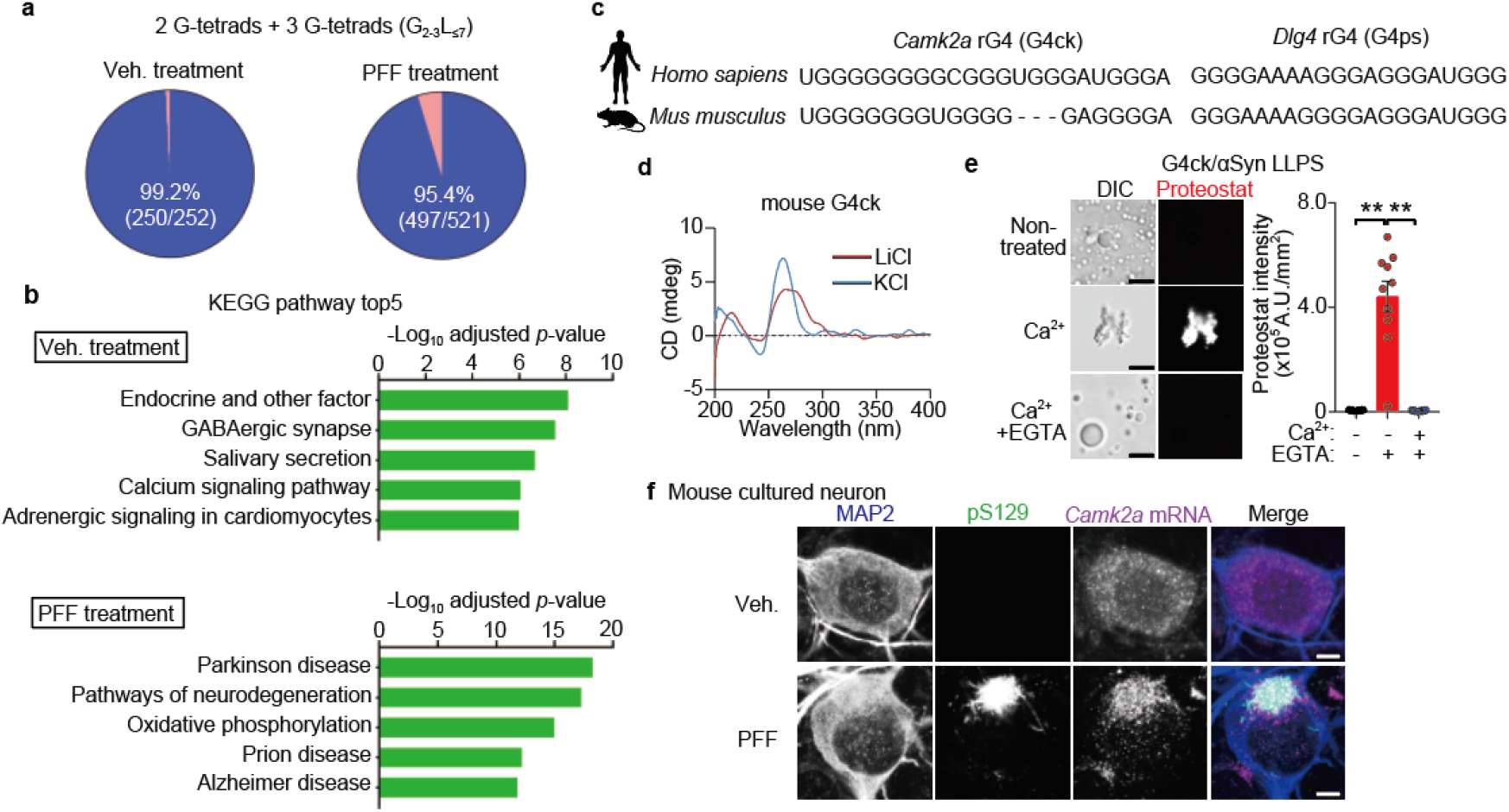
Bioinformatic and biophysical analyses of BG4-RIP-seq in pre-formed fibrils (PFF)-treated mouse cultured neurons. **a**, RNA G-quadruplexes (rG4s) in BG4-enriched RNAs containing two or three G-tetrads (G_2–3_L_≤7_) predicted by quadruplex-forming guanine-rich sequences (QGRS) mapper in mouse cultured neurons after PFF treatment. **b**, Kyoto Encyclopedia of Genes and Genomes (KEGG) pathway analysis in BG4-enriched RNAs in mouse cultured neurons after PFF treatment. **c**,**d**, Predicted rG4 sequences in human/mouse G4ck and G4ps (**c**) and circular dichroism spectra of mouse G4ck in the presence of KCl or LiCl (**d**). **e**, Representative images (left) and quantification (right) of Proteostat intensity within phase-transitioned G4ck/αSyn dependent on Ca^2+^. Scale bars, 5 μm. **f**, Representative images of MAP2 (blue), pS129 (green), and *Camk2a* RNA (magenta) in mouse cultured neurons after PFF treatment. Scale bars, 5 μm. Data are presented as mean ± standard error of mean. \*\**P* < 0.01 by one-way analysis of variance with Bonferroni’s multiple comparisons test. Number of replicates is shown in Supplementary Table 9.

**Extended Data Fig. 8:**
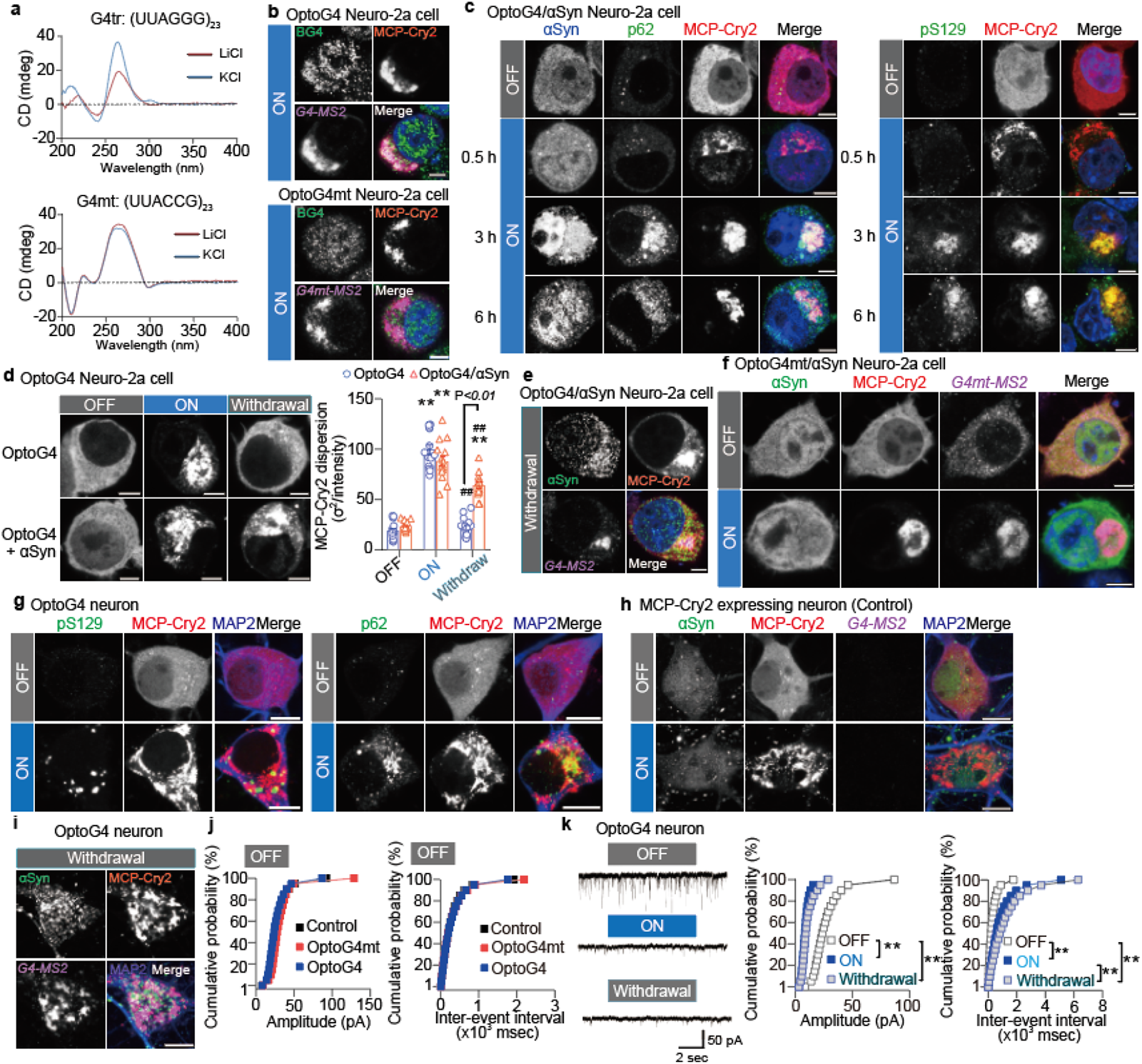
Establishment of forced RNA assembly in cells using optogenetics. **a**, Circular dichroism spectra of G4tr (UUAGGG)_23_ (left) and G4mt (UUACCG)_23_ (right) in the presence of KCl or LiCl. **b**, Representative images of G4tr (top) or G4mt (bottom) assembly in OptoG4-(top) and OptoG4mt-(bottom) expressing Neuro-2a cells respectively, with 3-h BL stimulation. Scale bars, 5 μm. **c**, Representative images of p62 (left) and pS129 (right) in Neuro-2a cells co-expressing optoG4 and α-synuclein (αSyn) with blue light (BL) stimulation. Scale bars, 5 μm. **d**, Representative images (left) and quantification (right) of MCP-Cry2 dispersion in Neuro-2a cells co-expressing optoG4 and αSyn for 3 hours of BL stimulation followed by 3 hours of withdrawal. Scale bars, 5 μm. **e**, Representative images of co-assembly of αSyn (green) and *G4-MS2* RNA (magenta) in Neuro-2a cells co-expressing optoG4 and αSyn under the withdrawal condition. Scale bar, 5 μm. **f**, Representative images of Neuro-2a cells co-expressing optoG4mt and αSyn with 3-h BL stimulation. Scale bars, 5 μm. **g**, Representative images of pS129 (left) and p62 (right) in optoG4-expressing mouse cultured neurons with 3-h BL stimulation. Scale bars, 5 μm. **h**, Representative images in MCP-Cry2-expressing mouse cultured neurons with BL stimulation for 3 h. Scale bars, 5 μm. **i**, Representative images of αSyn (green), MCP-Cry2 (red), *G4*-*MS2* RNA (magenta), and MAP2 (blue) in optoG4-expressing mouse cultured neurons under the withdrawal condition. Scale bar, 5 μm. **j**, Cumulative probability of sEPSCs related to Fig. 4f without BL stimulation. **k**, sEPSCs in OptoG4-expressing mouse cultured neurons for 3 hours of BL stimulation followed by 3 hours of withdrawal. Data are presented as mean ± standard error of mean. \*\**P* < 0.01 vs. non-stimulated group and ^##^*P* < 0.01 vs. stimulated group by one-way analysis of variance (ANOVA) with Bonferroni’s multiple comparisons test (**d**). \*\**P* < 0.01 by two-way ANOVA with Bonferroni’s multiple comparisons test (**k**). Number of replicates is shown in Supplementary Table 9.

**Extended Data Fig. 9:**
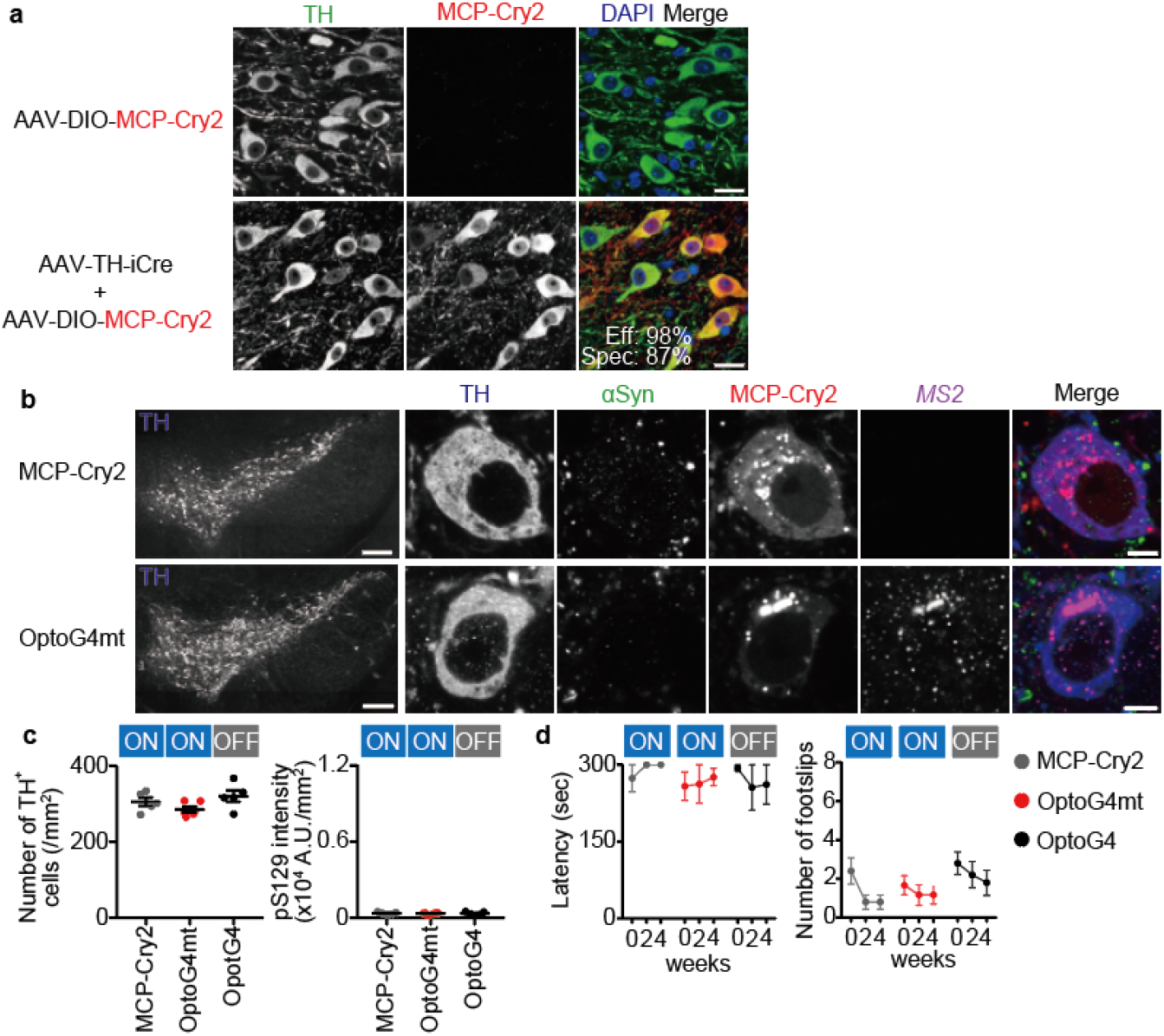
Forced RNA assembly in mouse nigral dopaminergic neurons by optogenetics. **a**, Representative images of tyrosine hydroxylase (TH) (green), MCP-Cry2 (red), and 4’,6-diamidino-2-phenylindole (blue) in the substantia nigra two weeks after the injection of AAV-DIO-MCP-Cry2 with AAV-pTH-iCre. Eff, efficiency; Spec, specificity. Scale bars, 20 μm. **b**, Representative images of TH (blue), αSyn (green), MCP-Cry2 (red), and *MS2* RNA (magenta) in the substantia nigra with blue light (BL) stimulation. Low and high magnification scale bars, 200 and 5 μm, respectively. **c**, Quantification of the number of TH-positive cells (left) and the fluorescent intensity of pS129 (right) in the substantia nigra with BL stimulation. **d**, Motor function in rotarod (left) and beam-walking (right) tasks with BL stimulation measured at the indicated time. Data are presented as mean ± standard error of mean. Number of replicates is shown in Supplementary Table 9.

**Extended Data Fig. 10:**
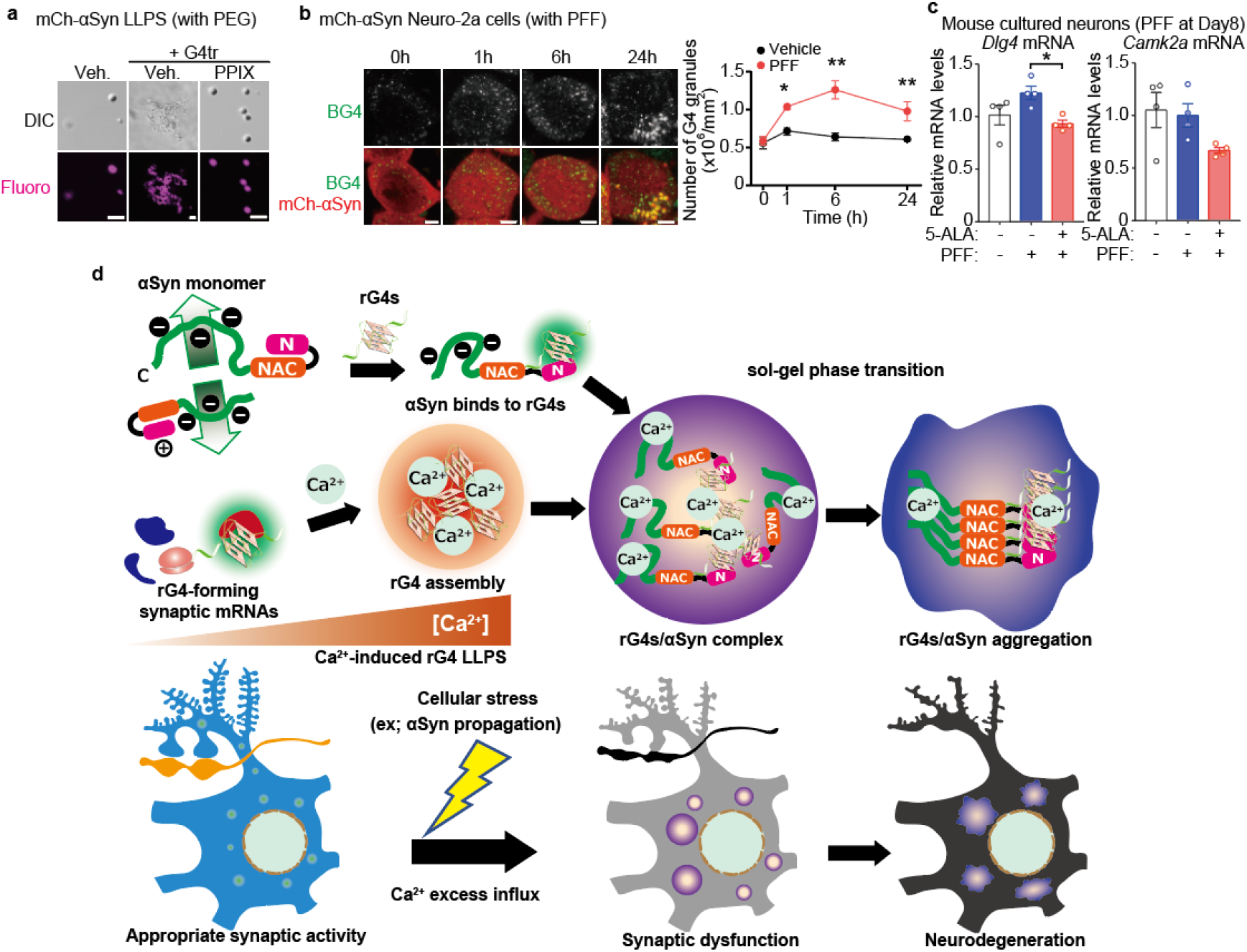
Effects of G4 ligand on RNA G-quadruplexes (rG4)-induced α-synuclein (αSyn) aggregation. **a**, Representative images of *in vitro* phase-transitioned G4tr/αSyn dependent on polyethylene glycol) and the effect of protoporphyrin IX. Scale bars, 5 μm. **b**, Representative images (left) and quantification (right) of BG4 granules (green) of Neuro-2a cells expressing mCh-αSyn (red) followed by pre-formed fibril (PFF) treatment for the indicated time. Scale bars, 5 μm. **c**, *Dlg4* and *Camk2a* mRNA levels in mouse cultured neurons following PFF treatment and the effects of 3 μM 5-aminolevulinic acid (5-ALA). **d**, Schematic showing rG4 assembly-induced αSyn aggregation and neurodegeneration: The monomeric state of αSyn is maintained by biased electrostatic and hydrophobic intramolecular interactions and intermolecular repulsion. rG4s bind to N-terminus of αSyn directly and then promote its aggregation. In the presence of Ca^2+^, enhanced rG4 assembly via liquid-liquid phase separation may serve as scaffolds for αSyn aggregation contributing to neurodegeneration. When neurons suffer from cellular stress such as αSyn propagation, αSyn aggregates with rG4 assembly composed of G4-forming synaptic mRNAs by Ca^2+^ excess influx, inhibiting synaptic protein translation, eliciting synaptic dysfunction, and eventually resulting in neurodegeneration. \**P* < 0.05 and \*\**P* < 0.01 by two-way (**b**) and one-way (**c**) analysis of variance with Bonferroni’s multiple comparisons test. Number of replicates is shown in Supplementary Table 9.

**Supplementary Table 1. Enriched 12-mer sequences in *α*Syn and BG4 by MERMADE program.**

**Supplementary Table 2. BG4-enriched mRNAs in vehicle-treated mouse cultured neurons.**

**Supplementary Table 3. BG4-enriched mRNAs in pre-formed fibrils-treated mouse cultured neurons.**

**Supplementary Table 4. Prediction of G4 structure in BG4-enriched mRNAs in vehicle-treated mouse cultured neurons by quadruplex-forming guanine-rich sequences mapper.**

**Supplementary Table 5. Prediction of G4 structure in BG4-enriched mRNAs in pre-formed fibrils-treated mouse cultured neurons by quadruplex-forming guanine-rich sequences mapper.**

**Supplementary Table 6. Kyoto Encyclopedia of Genes and Genomes (KEGG) pathway analysis in BG4-enriched mRNAs in vehicle-treated neurons.**

**Supplementary Table 7. Kyoto Encyclopedia of Genes and Genomes (KEGG) pathway analysis in BG4-enriched mRNAs in pre-formed fibrils-treated neurons.**

**Supplementary Table 8. Gene Ontology (GO) analysis in BG4-enriched mRNAs in pre-formed fibrils-treated neurons.**

**Supplementary Table 9. Statistical analysis and number of replicates. Supplementary Method.**

**Supplementary Video. Live imaging of pre-formed fibrils (PFF)-treated cells following GFP-*α*-synuclein (*α*Syn) expression in a fluorescence recovery after photobleaching (FRAP) assay**.

FRAP assay in HEK293T cells-expressing GFP-αSyn treated with PFF for the indicated time. Scale bars, 2 μm. Open circles in green indicate the regions of photobleaching.

