## Supplementary material for "RNA G-quadruplexes forming scaffolds for α-synuclein aggregation lead to progressive neurodegeneration": Supplementary Data.pdf

a

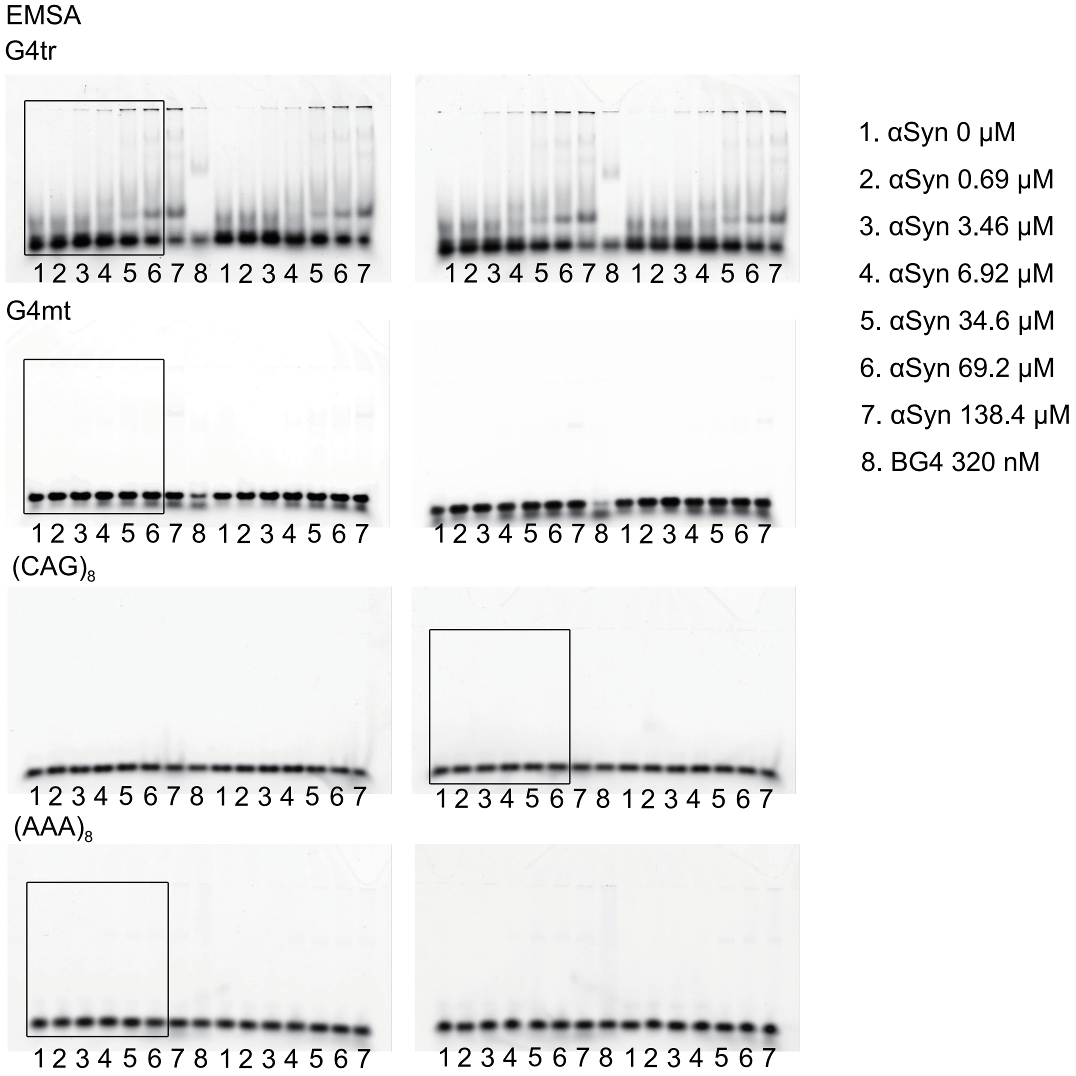


b

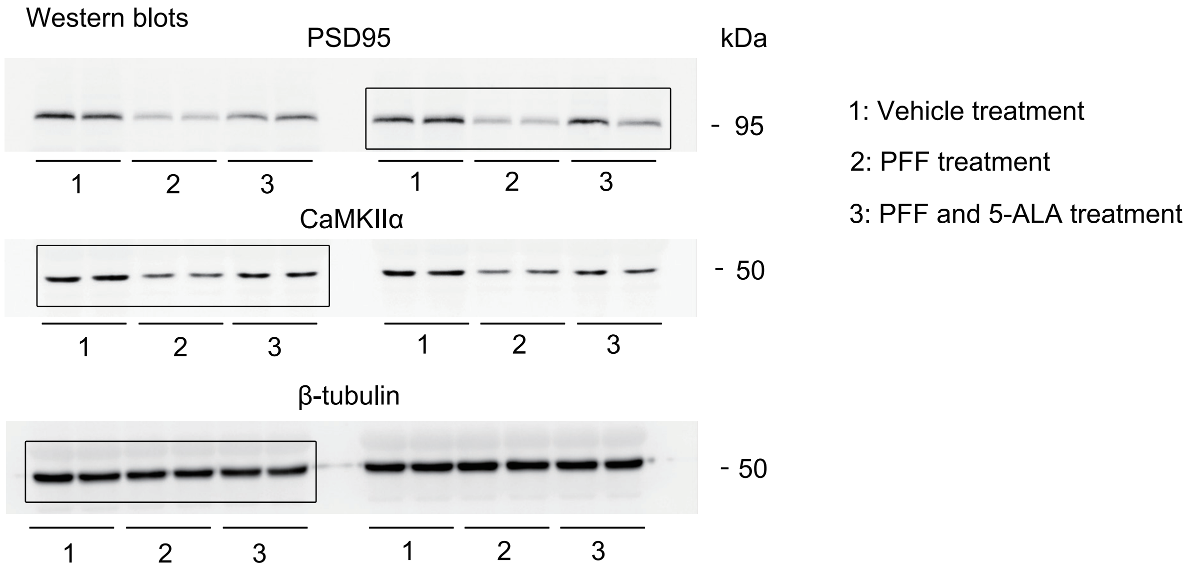


Supplementary Data 1. Full size scans of EMSA in Fig.1g and Extended Data Fig.4b and western blots shown in Fig.5f.
