## Supplementary material for "RNA G-quadruplexes forming scaffolds for α-synuclein aggregation lead to progressive neurodegeneration": Supplementary Method.docx

**Antibodies**

The following primary antibodies were used: anti-αSyn (1:1,000, ab212184, Abcam; 1:1,000, 4179S, Cell Signaling Technology), anti-pS129 (1:1,000, ab51253, Abcam), anti-DYKDDDDK tag (1:800, 14793S, Cell Signaling Technology; 1:10,000, KO602-L, Medicinal Chemistry Pharmaceutical), anti-6×His epitope tag (1:2,000, ab18184, Abcam), BG4 (1:800) (Asamitsu S et al., 2020), anti-Digoxigenin (1:1,000, ab76907, Abcam), anti-MAP2 (1:10,000, ab5392, Abcam), anti-p62 (1:1,000, GP62-C, Progen), anti-FMR1 (1:200, sc101048, Santa Cruz Biotechnology), anti-DCP1a (1:100, sc100706, Santa Cruz Biotechnology), anti-HuR (1:100, sc5261, Santa Cruz Biotechnology), anti-TH (1:1,000, ab76442, Abcam; 1:1,000, 22941, Immunostar), anti-PSD95 (1:1,000, 3450S, Cell Signaling Technology), anti-CaMKIIα (1:1,000, 50049S, Cell Signaling Technology), and anti-β-tubulin (1:5,000; T8328, Sigma-Aldrich). For immunoblotting, the following secondary antibodies were used: horseradish peroxidase (HRP)-conjugated anti-mouse (1:5,000, 1031-05, SouthernBiotech) and HRP-conjugated anti-rabbit (1:5,000, 4050-05, SouthernBiotech). For immunohistochemistry, the following secondary antibodies were used: DyLight 405-AffiniPure donkey anti-rabbit (1:1000, 715-475-150, Jackson ImmunoResearch), Alexa 488-conjugated donkey anti-mouse (1:1,000, A-21202, Invitrogen), Alexa 594-conjugated donkey anti-mouse (1:1,000, A-21203, Invitrogen), DyLight 405-AffiniPure donkey anti-rabbit (1:1,000, 711-475-152, Jackson ImmunoResearch), Alexa 488-conjugated donkey anti-rabbit (1:1,000, A-21206, Invitrogen), Alexa 594-conjugated donkey anti-rabbit (1:1,000, A-21207, Invitrogen), Alexa 594-AffiniPure Donkey Anti-Guinea Pig (1:1,000, 706-585-148, Jackson ImmunoResearch), Cy5-AffiniPure Donkey Anti-Goat (1:1,000, 705-175-147, Jackson ImmunoResearch), and DyLight 405-AffiniPure donkey anti-chicken IgY (1:1,000; 703-475-155, Jackson ImmunoResearch).

**FISH probes**

The following 5'-DIG-labeled probes were synthesized by Integrated DNA Technologies and NIHON GENE RESEARCH LABORATORIES: *MS2* (5'-TTTCTAGAGTCGACCTGCAG-3', 5'-CTAGGCAATTAGGTACCTTAG-3', and 5'-CTAATGAACCCGGGAATACTG-3')^55^, *Camk2* (5'-TGCGTCCAAGTAGAAGTGATACCTAAATGTTGCTTGCTTTGC-3')^56^, and *Dlg4* (5'-AGGGGGCGTGTCTTCATCTTGGTAGCGGTATTTCTTGGTTGTCAC-3')^56^.

**RT-qPCR primers**

Sample preparation for RT-qPCR from primary cultured neurons was performed using the RNeasy Mini Kit and PrimeScript RT Master Mix (Takara Bio). RT-qPCR was performed using KOD SYBR qPCR Mix (TOYOBO) on a CFX Connect Real-Time PCR System (Bio-Rad Laboratories). Gene expression was assessed using the differences in normalized Ct (cycle threshold) (^ΔΔ^Ct) method after normalization to *Gapdh* expression. Fold change was calculated by 2^-ΔΔCt^. The following primers were used for RT-qPCR: *Camk2a* (forward, 5'-AGCCATCCTCACCACTATGCTG-3'; reverse, 5'- GTGTCTTCGTCCTCAATGGTGG-3') (OriGene), *Dlg4* (forward, 5'-GCCCTGTTTGACTACGACAA-3'; reverse, 5'-CTCATAGCTCAGAACCGAGT-3')^57^, and *Gapdh* (forward, 5'-AACTTTGGCATTGTGGAAGG-3'; reverse, 5'-ACACATTGGGGGTAGGAACA-3').
